# Traceless protein semi-synthesis in cells using the promiscuous ultra-fast split intein NrdJ-1

**DOI:** 10.64898/2026.03.03.709324

**Authors:** Xuanjia Ye, Joshua Sokol, Christian Kofoed, Anushka Dheer, Juner Zhang, Tom W. Muir

## Abstract

Chemistry-driven approaches that allow proteins to be manipulated in ways not permitted by standard genetics are indispensable in the basic and applied biomedical sciences. Of the various platform technologies in common use, those that exploit intein-mediated protein splicing have proven especially powerful owing to compatibility with both *in vitro* and *in vivo* applications. Here, we provide detailed biochemical characterization of the split intein, NrdJ-1. We demonstrate that the rapid splicing kinetics associated with this system are largely independent of splice junction primary sequence context. The promiscuity and high efficiency of this split intein makes it well suited to the generation of chemically modified proteins *in vitro* and in living cells, as demonstrated by the traceless generation of semisynthetic chromatin harboring various post-translational modifications. These studies highlight the utility of NrdJ-1 for traceless protein editing and suggest it is a superior choice for many intracellular biochemical applications.

## Introduction

Understanding the biochemical roles of post translational modifications (PTMs) remains an extremely challenging undertaking^1–3^. Unambiguous assignment of function requires the ability to introduce a given PTM site-specifically into an otherwise native protein sequence, something that cannot be achieved using standard genetic approaches. Several complementary chemistry-driven methods have been developed in recent decades that allow the incorporation of a wide range of non-canonical amino acids (ncAAs), including many PTMs, into proteins^4–6^. Approaches such as genetic code expansion, chemical protein (semi)synthesis and sidechain-directed conjugations have become important weapons in the biomedical research arsenal, allowing incorporation of biochemical probes and PTMs into those proteins accessible to the methods. Among these approaches, protein semisynthesis, a strategy involving the assembly of recombinant with chemically synthesized polypeptide fragments, has emerged as especially powerful for the study of PTMs due to the range of chemotypes that can be introduced at one or more discrete positions in a protein^4^.

While protein semisynthesis is widely used in the *in vitro* (herein defined as test-tube biochemistry) analysis of PTMs – for example, we and others have introduced numerous modifications into histone proteins for biochemical studies^7^ – methods such as native chemical ligation (NCL) and expressed protein ligation (EPL) cannot be directly implemented in cells due to the requirement for high reactant concentrations and the instability of reaction intermediates under physiological conditions^8,9^. By contrast, chemo-enzymatic strategies offer greater flexibility in terms of compatibility with both *in vitro* and cellular applications^4^. Among these, split intein-mediated protein trans-splicing (PTS) is especially attractive due to the high specificity and autocatalytic nature of the process ^4,10,11^. In PTS, complementary N- and C-terminal split inteins (designated Int^N^ and Int^C^) associate and mediate the ligation of the polypeptides they are fused to which, by convention, are referred to as the exteins. If one of the extein sequences is a synthetic peptide then the resulting PTS reaction results in the semisynthesis of a protein (Figure 1). Split intein-mediated PTS have been used in numerous protein engineering applications *in vitro*, in living cells and *in vivo*^10,12,13^.

**Figure 1.**
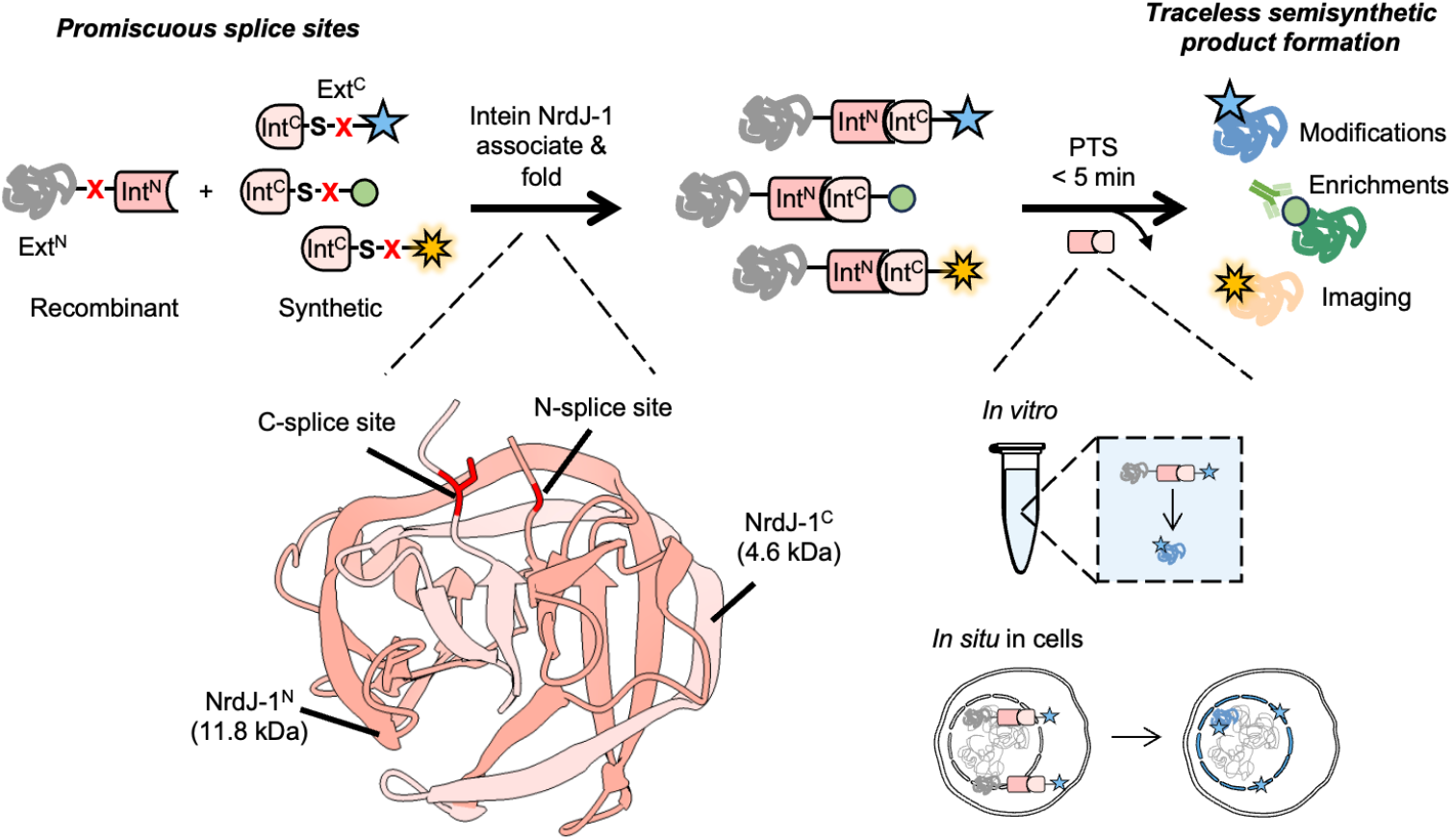
Schematic of NrdJ-1 mediated protein trans-splicing (PTS), featuring promiscuous splice sites and traceless, ultrafast ligation both *in vitro* and in cells. NrdJ-1 fragments (Int^N^, Int^C^) spontaneously associate and fold to initiate ligation of the adjoining exteins upon mixing. Minimally, a serine residue is required at the splice junction. For protein semisynthesis, a synthetic fragment bearing user-defined chemical functionalities (indicated by colored stars) is ligated to a recombinant protein of interest. This approach enables installation of diverse chemical modifications, enrichment handles, or imaging probes and can be performed either *in vitro* or *in situ* within the native cellular environment. Structure of the folded NrdJ-1 intein is shown (PDB: 8UBS).

Since PTS involves an excision-ligation reaction, the process has the potential to return a final protein product with few if any alterations to the native primary sequence. This differentiates the approach from the commonly used enzymatic ligation method based on bacterial sortases (i.e. sortagging) which requires short peptide sequences in both reactants and thus typically leaves behind a ‘ligation scar’ in the product^14^. In practice, inteins have evolved to efficiently splice in a native extein context, meaning that they often have local sequence preferences either side of the splice junction^15,16^. Moreover, PTS minimally requires a Cys, Ser or Thr residue immediately adjacent to the C-intein fragment which places further constraints on the choice of ligation sites in a target protein sequence^4^. Deviations from these optimal sequences can perturb the alignment of key residues in the intein splicing center leading to the accumulation of side-products and slower splicing kinetics^16^. Even in an optimal extein context, different split inteins support PTS with varied kinetics. In the best cases, the half-life for PTS can be on the order of seconds, making these ‘ultrafast’ split inteins particularly appealing for protein engineering applications^4^.

Of the ultrafast split inteins reported to date, the NrdJ-1 system, first identified in a marine metagenomic study, is especially attractive for protein semisynthesis^17–22^. While most characterized ultrafast split inteins require a cysteine at the first C-extein position, NrdJ-1 utilizes a serine (Figure S1), thereby offering expanded flexibility in selecting splice sites due to the substantially higher abundance of serine (∼8–9%) compared to cysteine (∼2–3%) in the human proteome^23^. Moreover, several recent studies have hinted that NrdJ-1 might have relaxed extein dependence^21,22,24^, although a systematic evaluation of this is yet to be reported. In this study, we provide a detailed characterization of NrdJ-1, including an assessment of its local extein sequence tolerance. These studies reveal NrdJ-I to be a remarkably promiscuous split intein and provide insights into the underlying basis of its ultrafast splicing kinetics. Motivated by these studies, we show that this split intein can be used for traceless protein semisynthesis *in vitro* and in cells, with a particular focus on editing the highly modified N-terminal tail of histone H3.

## Result

### NrdJ-1 exhibits low extein dependence

To explore the extein dependence of NrdJ-1, we first generated a series of recombinant protein constructs of the general structure, MBP–X_-2_X_-1_–NrdJ-1^N^ and NrdJ-1^C^– X_1_X_2_X_3_–eGFP, where MBP and eGFP are model N- and C-extein proteins, respectively, and X_n_ denotes a variable amino acid residue on either side of the splicing junction (Figure 2A). We focused on four positions known to influence intein activity: –1 and –2 on the N-extein, and +2 and +3 on the C-extein^15,16,19,25^. Note, the +1 position in the C-extein was kept constant as a serine, since this is required for NrdJ-1 mediated PTS. Following bacterial expression, these proteins were purified and characterized by LC-MS (Figure S2). As a benchmark, we first determined the splicing rate of NrdJ-1 in a native extein context. The requisite proteins were combined (1 μM each) at 37 °C and the reaction progress was monitored over time by SDS-PAGE and LC-MS (Figure 2B, Figure S2A and Figure S4A, Supplementary table 1). Under these conditions, NrdJ-1 supported rapid and highly efficient splicing with an apparent first order rate constant *k*_splice_ = 0.027 ± 0.001 s^-1^, corresponding to a t_½_ = 26.0 ± 1.2s, confirming its ultrafast kinetics and high catalytic efficiency under physiological conditions^17^.

**Figure 2.**
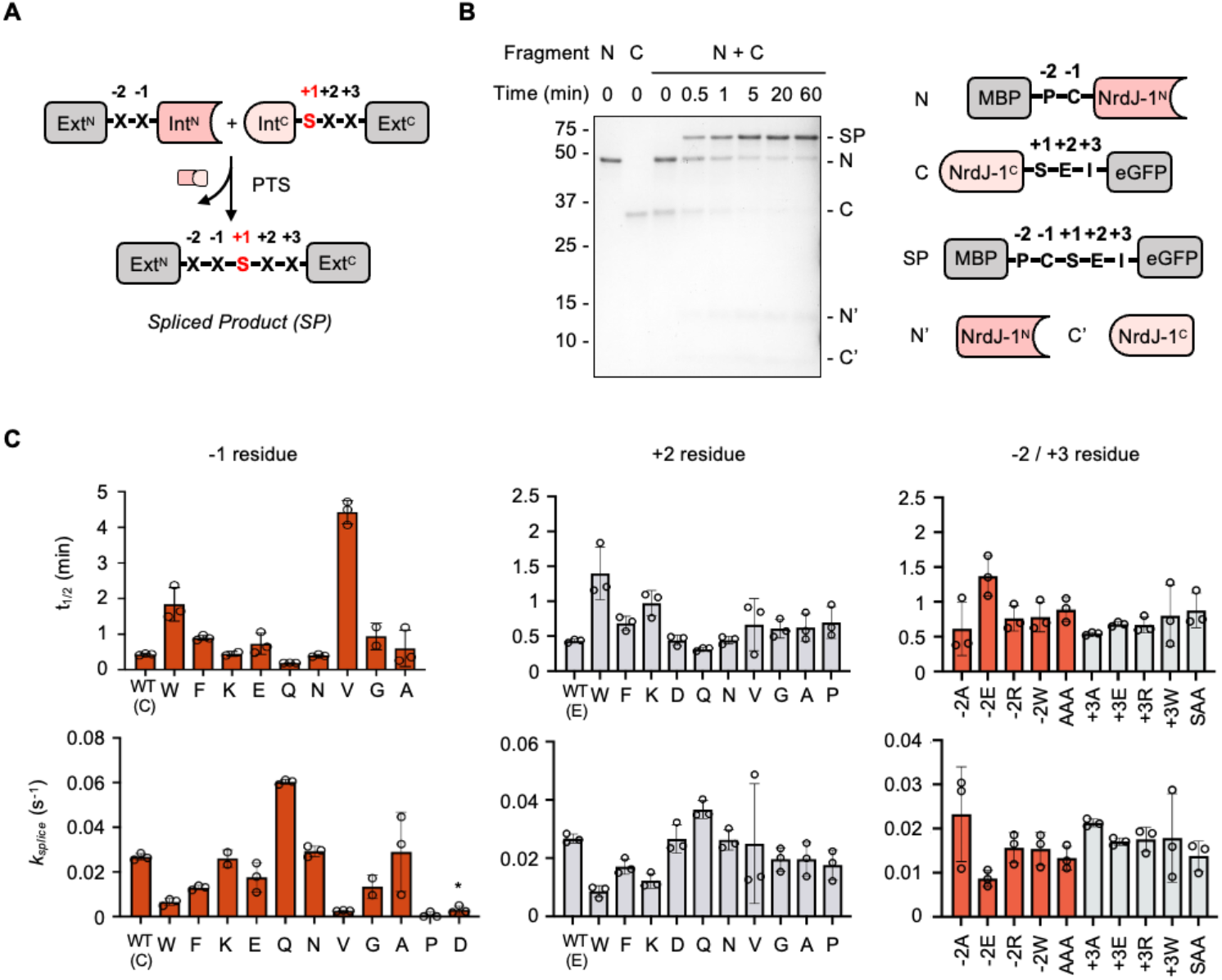
NrdJ-1 exhibits low extein dependence. (A) Schematic showing the splicing junction of PTS. Ext^N/C^ and Int^N/C^ refer to the N- and C-terminal exteins and split inteins, respectively. Extein residues flanking the splice site are indicated. For NrdJ-1 the +1 position is an obligate serine (red). (B) Time course of NrdJ-1 PTS reaction featuring native extein residues at the splice junction. MBP-NrdJ-1^N^ and NrdJ-1^C^-eGFP were combined (1 μM of each reactant) in splicing buffer (pH 7.2) at 37°C and the reaction progress was monitored by SDS-PAGE with Coomassie staining. Identity of the spliced product (SP) was confirmed by mass spectrometry (see Figure S2). The schematic on the right depicts the starting materials and corresponding end products. (C) Impact of varying indicated N- and C-extein residues on the kinetics. Reactions were performed and monitored as in panel B and the extent of product determined by gel densitometry analysis (details in Material and Methods). The half-lives (upper panel) and apparent first order rate constants (lower panel) for the reactions are presented. * above -1 Asp indicates side product formation (see Figure S4). Error bars represent SD (n = 2-3).

Next, we assessed the splicing activities of the constructs containing mutations in the variable positions at the splice junction (Figure 2C and Figure S3, Supplementary table 1). The mutations were designed to explore the impact of altering the charge (e.g. Lys, Glu, Asp), size (e.g. Ala, vs. Trp/Phe) and conformation properties (Pro, Gly) of the residues on splicing efficiency. These studies revealed NrdJ-1 possesses remarkable extein tolerance, with high spliced product yields and fast kinetics observed for most mutants explored. The exceptions to this were Asp and Pro at the -1 position which significantly compromised reaction fidelity (Figure 2C, Figure S3 and Figure S4B, C). Both these residues are well known to be incompatible with intein-mediated protein splicing^4,16^; Asp promotes premature N-extein cleavage through a cyclization/hydrolysis process (Figure S4C-F), whereas the presence of a - 1 Pro disrupts the first step in splicing involving an N-to-S(O) acyl shift. Interestingly, Pro was found to be compatible when placed at the +2 extein position (Figure 2C and Figure S3B), which is not the case for many other ultrafast inteins such as the commonly used DnaE family of split inteins^25^. Further underlining the flexibility of the NrdJ-1 system, we found that efficient splicing was supported when multiple mutations were made at the N-(AAA) and C-splice junctions (SAA) (Figure 2C and Figure S3C). Together these studies reveal NrdJ-1 to have exceptionally low extein-dependence, making it an excellent candidate for further engineering and applications.

### Mechanisms of NrdJ-1 rapid kinetics

We next turned to the underlying basis of the fast splicing kinetics associated with NrdJ-1. The first step in PTS is the association of the Int^N^ and Int^C^ fragments (Figure S1). Naturally split inteins often exhibit high affinity between the fragments, a property that is desirable for cellular applications where low concentrations of the reactants are present^26^. Since the binding affinity of NrdJ-1 fragments has not previously been reported, we performed isothermal titration calorimetry (ITC) using MBP–NrdJ-1^N^ and NrdJ-1^C^–SUMO constructs containing mutations that block splicing chemistry but not fragment association (C1A in the N-intein and N145A in the C-intein, Figure S5A, S5B). These fusion partners were selected to mimic the interaction environment of NrdJ-1 when appended to proteins in cellular applications, rather than measuring the affinity of the isolated intein fragments alone. For this experiment, native NPC and SEI extein sequences were retained at the splice junctions. This analysis revealed tight association between the fragments, with ∼1:1 stoichiometry and a dissociation constant (K_d_) of 68.4 ± 5.6 nM (Figure 3A and Figure S5C).

**Figure 3.**
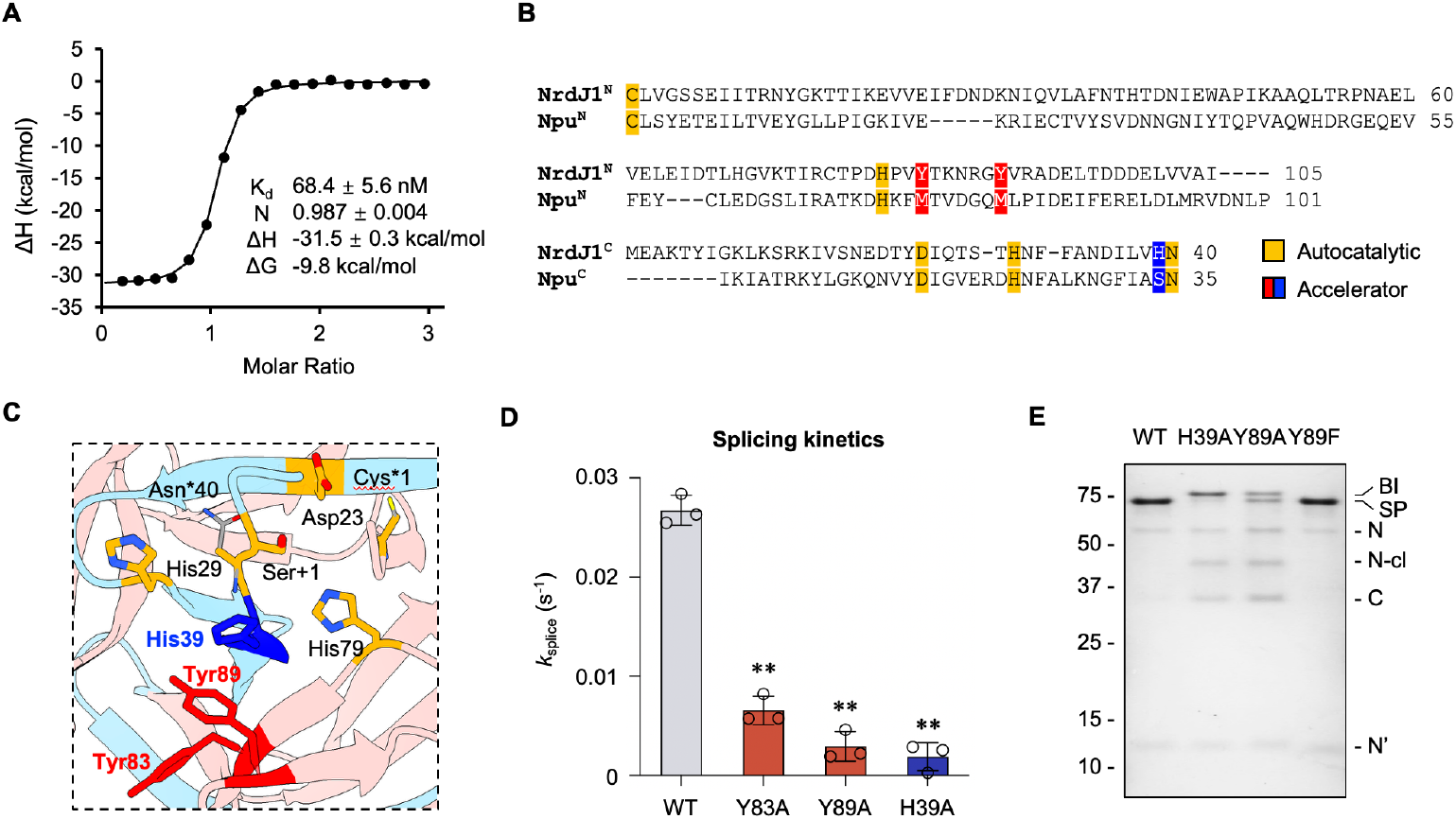
‘Accelerator’ residues facilitate succinimide formation in the splicing reaction. (A) Thermogram and thermodynamic parameters of NrdJ-1 fragment association. Analysis was performed by ITC using MBP-NrdJ-1^N^ and NrdJ-1^C^-SUMO bearing inactivating mutations (C1A and N145A). K_d_: dissociation constant; n: stoichiometry; ΔH: enthalpy change; ΔG: Gibbs free energy change. (B) Multiple sequence alignment of the intein fragments of NrdJ-1 with a representative fast-splicing split intein, Npu. Conserved residues involved in autocatalysis and identified accelerators are highlighted. (C) Structure of the NrdJ-1 split intein showing the spatial positions of conserved autocatalytic residues (Cys1, Asp23, His29, Asn40, Ser+1, highlighted in orange) and the candidate accelerator residues (Tyr83, Tyr89 within NrdJ-1^N^ and highlighted in red and His39 within NrdJ-1^C^ and colored blue). The regions corresponding to the N- and C-intein fragments in the split version are colored pink and blue, respectively. The structure is generated from an engineered fused version of NrdJ-1 bearing inactivating mutations: C1A and N145A. The side chains of Cys1 and Asn145 were modeled in silico using Chimera rotamer simulations. PDB: 8UBS. (D) *In vitro* PTS kinetics with wildtype NrdJ-1 and corresponding accelerator mutants. *k*_*splice*_ represents first-order kinetic fits of splice product formation (see Materials and Methods). Reactions were carried out as in Figure 2 and products quantified from densitometry analysis of SDS-PAGE analyses of time courses. Error bar indicates SD (n = 3); **: p ≤ 0.01. (E) PTS reactions employing wild-type or indicated mutant PTS mixtures. Reactions were quenched at 30 mins and visualized by SDS-PAGE followed by Coomassie staining. BI: branched intermediate; SP: spliced product; N: MBP-NrdJ-1^N^; N-cl: N-terminal cleavage; C: NrdJ-1^C^-eGFP; N’: NrdJ-1^N^ after splicing.

The splicing activity of split inteins relies on a conserved network of autocatalytic residues organized within the HINT (Hedgehog/Intein) fold^10^ (Figure S6A). In addition to this well-defined autocatalytic core, previous studies on the DnaE family of split inteins have identified a class of auxiliary positions, referred to as “accelerator” residues, that enhance splicing efficiency^27^. These residues are located in the second coordination shell relative to the autocatalytic core and have been shown to promote rapid splicing by stabilizing key transition states or intermediates. To identify potential “accelerator” residues in NrdJ-1, we performed a multiple sequence alignment (MSA) with the well-characterized ultrafast DnaE intein, Npu^27^. As expected, this revealed strong conservation of the canonical autocatalytic residues, consistent with the shared structural homology of the two split inteins (Figure 3B and Figure S6B). Given this similarity, we hypothesized that NrdJ-1 might possess functional counterparts to the accelerator residues previously identified in Npu. Intriguingly, at the positions corresponding to the Met and Ser accelerator residues in Npu, NrdJ-1 instead contains aromatic residues. Structural mapping revealed that these residues lie adjacent to the autocatalytic core and occupy spatially analogous positions to the established accelerator sites in Npu (Figure 3C and S6B). To test their functional roles, we introduced alanine substitutions at Tyr83 and Tyr89 in the NrdJ-1^N^ fragment and His39 in NrdJ-1^C^ (Figure S7A). All three mutants displayed significant reductions in splicing kinetics relative to the wildtype intein, indicating that these residues contribute to efficient autocatalysis (Figure 3D and Figure S8A, B). Consistent with their spatial proximity to the autocatalytic core, alanine mutations at His39 and Tyr89 exerted a stronger impact on splicing efficiency than the Y83A mutation (Figure 3C, D, and Figure S8A, B).

To probe the mechanistic role of the putative “accelerator” residues, we further analyzed the reaction products obtained when using alanine mutants. SDS-PAGE analysis of the reaction mixtures revealed that the H39A and Y89A variants accumulated a distinct high molecular-weight species not observed in the wildtype intein (Figure 3E and Figure S8B). Subsequent analysis by RP-HPLC and mass spectrometry identified this species as the branched intermediate (BI) wherein the N-extein is linked to the +1 serine through an ester linkage (Figure S1A, Figure S7B and S8C). This intermediate is on the canonical reaction pathway for protein splicing and its resolution through succinimide formation is typically the rate-limiting step in the process^28^. The accumulation of BI suggests a defect in this step. Notably, replacement of Tyr89 with a phenylalanine did not result in BI accumulation (Figure 3E, Figure S8D, S8E), suggesting that the presence of an aromatic sidechain at this position is critical for the resolution of this key splicing intermediate. Inspection of the crystal structure of NrdJ-1 reveals that Tyr89 packs against both Tyr83 and His39 (Figure S6B). Importantly, His39 is immediately adjacent to Asn40 in NrdJ-1^C^, the residue that directly participates in succinimide formation. Previously, we have shown that even minor perturbations to the positioning of the Asn side chain perturbs this reaction^16,28^. Thus, it seems reasonable to propose that disruption of the Tyr83-Tyr89-His39 network would interfere with this step in the splicing reaction. More generally, tuning the activity of split inteins through manipulation of ‘accelerator residues’ was key to the recent development of a protein editing technology that involves the activity of matched pairs of orthogonal split inteins^29^. We expect that the above insights into the NrdJ-1 splicing are likely to be useful for its integration into this protein engineering manifold.

### Traceless serotonylation on histone H3 tails

The biochemical features of NrdJ-1, its efficiency and broad extein tolerance, motivated us to evaluate its utility across multiple contexts. First, we sought to perform *in vitro* protein semisynthesis involving installation of PTMs with the view to subsequently extend the workflow to cellular applications. Histone H3 represents an ideal target for such efforts: its N-terminal tail is densely modified with regulatory PTMs that influence chromatin structure and gene expression^2^. Importantly, the native H3 tail lacks cysteine, rendering cysteine-dependent PTS strategies non-traceless (Figure S9). Although cysteines can be introduced and later converted to alanine via radical desulfurization, this approach is incompatible with many ncAAs and several PTMs. By contrast, the first 35 residues of H3 contain three native serine residues (S10, S28, and S31) that can serve directly as splice junctions for PTM conjugation by PTS (Figure 4A and Figure S9). Thus, NrdJ-1 offers a means to achieve scarless ligation at native serine sites, eliminating the need for desulfurization and substantially expanding the positional and chemical flexibility of histone semisynthesis.

**Figure 4.**
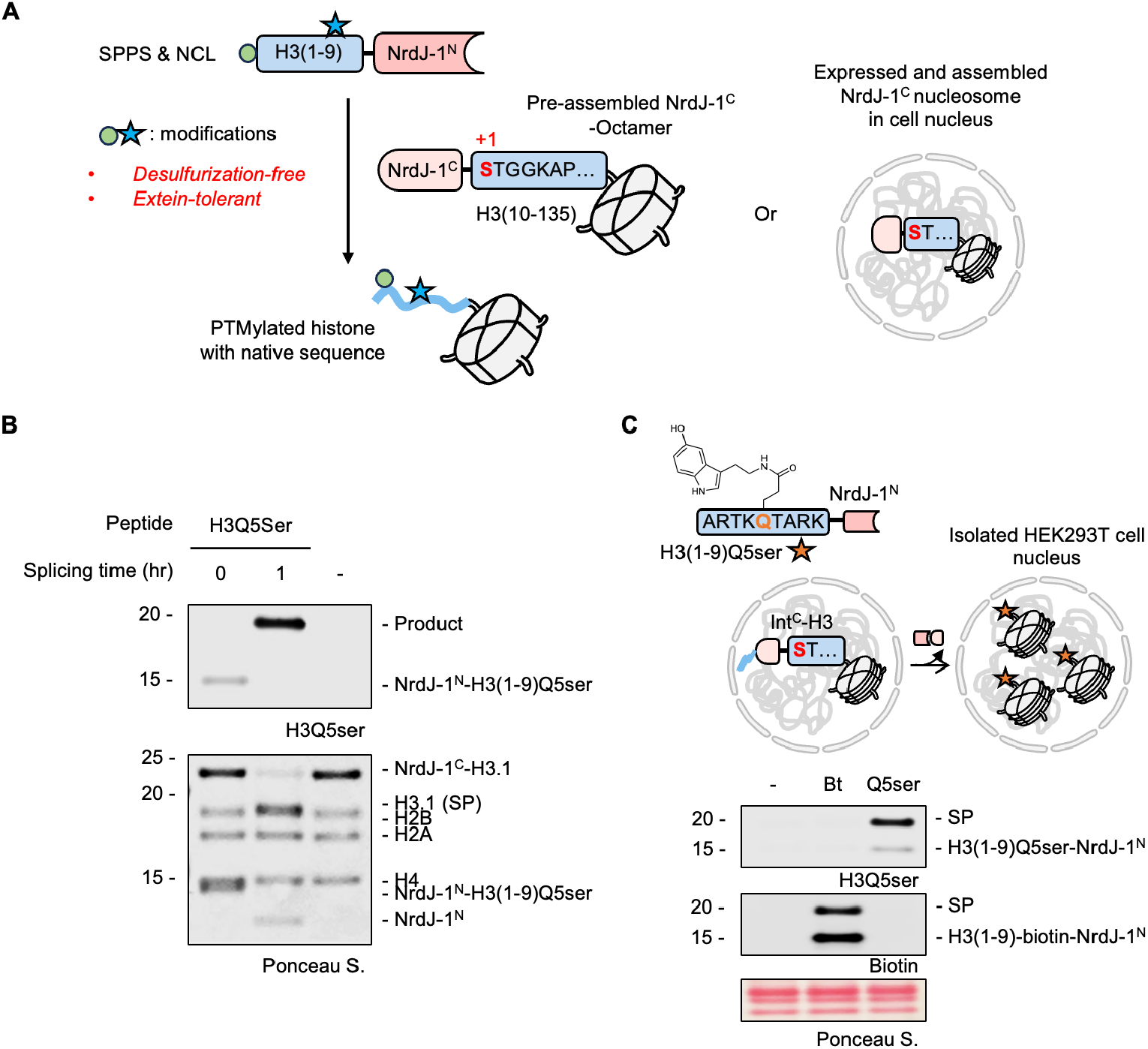
Traceless installation of H3Q5Ser into chromatin using NrdJ-1. (A) Schematic of histone H3 semisynthesis using NrdJ-1-mediated PTS. NrdJ-1^N^ fused to a synthetic PTM-bearing H3 peptide fragment (in the example shown residues 1-9) is reacted with the remainder of H3 (residues 10-135) fused to the complementary NrdJ-1^C^ intein. PTS leads to the traceless semisynthesis of modified H3, exploiting the ability of NrdJ-1 to utilize native serine residues. In principle, this workflow can either work in vitro on pre-assembled histone octamers or in isolated cell nuclei containing endogenously expressed chromatin-associated NrdJ-1^C^ histone fusion. (B) *In vitro* PTS reaction employing H3(1–9, Q5ser)-NrdJ-1^N^ and NrdJ-1^C^-H3(10–135) containing histone octamers. Reaction mixture was analyzed at indicated timepoint by western blot (top) and SDS-PAGE (bottom). (C) Top: schematic of the experimental design. The chemically modified substrate H3(1–9)-NrdJ-1^N^ containing either a biotin handle (Bt, serve as splicing control) or Q5ser was incubated with isolated HEK293T nuclei transiently expressing Flag-H3(1–28)-NrdJ-1^C^-H3(10–135). Protein trans-splicing excises the N-terminal Flag-H3(1–28)-NrdJ-1^C^ and ligates the modified H3(1–9) fragment to H3(10–135), installing biotin or Q5ser on chromatin; Bottom: western blot analysis of nuclear extracts after 1 h incubation at 37 °C. SP, spliced product.

Protein serotonylation involves the covalent attachment of serotonin, an essential neurotransmitter and hormone, to glutamine sidechains and is mediated by the enzyme transglutaminase 2 (TG2)^30^. Although discovered several decades ago, the diverse biological roles of protein serotonylation have only recently begun to emerge^31–34^. In particular, histone serotonylation at H3Q5 (H3Q5ser) is notably enriched in the gut and brain and has been implicated in establishing a permissive transcriptional state in serotonergic neurons^32^. While access to homogeneous preparations of H3Q5ser would greatly facilitate ongoing studies into the precise biochemical functions of this PTM, the incompatibility of serotonin with standard NCL/desulfurization workflows prevents the generation of truly authentic versions of this modified protein^31^. In principle, NrdJ-1-mediated semisynthesis provides a solution to this problem since PTS can exploit a native serine in the H3 N-terminal tail as a potential splice junction, thereby obviating the need for the problematic desulfurization step.

While histone H3 contains multiple native serine residues, we selected Ser10 as the splice junction for synthetic convenience (Figure 4A and S9). H3(1–9) peptides bearing Q5ser were prepared by solid phase peptide synthesis (SPPS), converted to thioesters, and ligated to NrdJ-1^N^ via NCL (Figure S10). The cysteine at position 76 in NrdJ-1^N^, previously shown to be dispensable for catalysis^22^ (Figure S11), was mutated to alanine to minimize disulfide formation; this variant was used for all subsequent semisynthesis unless otherwise noted. Rather than performing the PTS reaction on the isolated H3 fragments, we decided to explore whether the reaction could be performed at the level of histone octamers since this would, in principle, streamline the manufacture of chromatin for downstream *in vitro* studies. The purified NrdJ-1^C^-H3(10-135) construct was combined with recombinant histones H2A, H2B and H4 and then folded into octamers using established protocols (Figure S12A-C). Combining these purified histone octamers with H3(1-9, Q5ser)-NrdJ-1^N^ led to the efficient generation of the desired modified histone H3 as indicated by western blotting and LC-MS (Figure 4B and S12D). The success of the reaction is especially notable since the splicing buffer it was performed in contained 2 M NaCl, which is required for the stability of the histone octamer complex in the absence of DNA. The tolerance of the NrdJ-1 to such conditions further underlines the utility of this split intein system for protein semisynthesis.

To explore whether NrdJ-1–mediated histone editing could be conducted in a native chromatin setting, we expressed a histone H3.1 split intein fusion, Flag-H3(1-28)-NrdJ-1^C^–H3.1(10-135), in HEK293T cells. The H3(1-28) segment was included in this construct to enhance protein expression and chromatin incorporation^25^. Nuclei from these cells were isolated using established procedures^35^ and then treated with either a H3.1(1-9)-NrdJ-1^N^ fusion containing either an N-terminal biotin handle (Figure S13A), or the previously described H3(1-9, Q5ser)-NrdJ-1^N^ fusion. Following incubation for 1 h, the *in nucleo* PTS reactions were analyzed by western blotting (Figure 4C) which revealed the generation of the expected semisynthetic histones. Note, HEK293T cells do not contain endogenous H3Q5Ser levels, providing a clean background for assessing site-specific editing in this example. The success of the experiment represents the first traceless semisynthesis of a modified histone in a native chromatin context, an advance that will serve as a foundation for future studies gearing towards better understanding the biochemical functions of this PTM.

### Traceless histone H3 semisynthesis in live cells

Having established NrdJ-1-mediated installation of H3Q5 serotonylation *in vitro* and *in nucleo*, we next sought to (i) explore the scope of histone PTMs accessible to the system and (ii) apply this strategy in live cells to enable temporally controlled synthesis of PTM-modified histones. To this end, we selected three modifications, biotin, H3K9ac and H3K9me2, to evaluate the generality of the approach (Figure S13A-C). In addition to expanding beyond endogenous PTMs, biotin provides a versatile biochemical handle for visualization and affinity enrichment. H3K9me2 is an important mark for genome organization, whereas H3K9ac is associated with active transcription^36–38^; both modifications play key roles in chromatin regulation. Importantly, the lysine-9 residue occupies the −1 position relative to the splice junction, providing an additional testbed for extein tolerance (Figure S13D). To the best of our knowledge, PTS with a chemically modified −1 extein has not been previously demonstrated.

Consistent with our earlier findings, NrdJ-1 readily accommodated these sequence contexts both in pre-assembled octamers and in isolated nuclei expressing the NrdJ-1^C^-H3.1(10-135) fusion (Figure 5A and Figure S14). These results further demonstrate the flexibility of NrdJ-1 with respect to the choice of residues at the splice junction. We next sought to extend this approach to living cells. To deliver the synthetic intein fragments, we adopted a previously reported bead-loading method in which transient mechanical disruption of the cell membrane enables uptake of macromolecules (Figure S15A)^39,40^. We estimate that ∼10-15% of cells take up an exogenously added eGFP protein following bead loading (Figure S15B). As an initial test of this peptide delivery approach, we transiently expressed an mCherry-Lamin A-NrdJ-1^N^ construct in HEK293T cells and then introduced purified NrdJ-1^C^–eGFP via bead-loading (Figure S15C). Gratifyingly, we observed localized GFP signal at the nuclear periphery only in those cells expressing the Lamin A-NrdJ-1^N^ fusion, whereas no GFP signal was detected in the negative controls without protein or bead loading processing. (Figure S15D and S15E).

**Figure 5.**
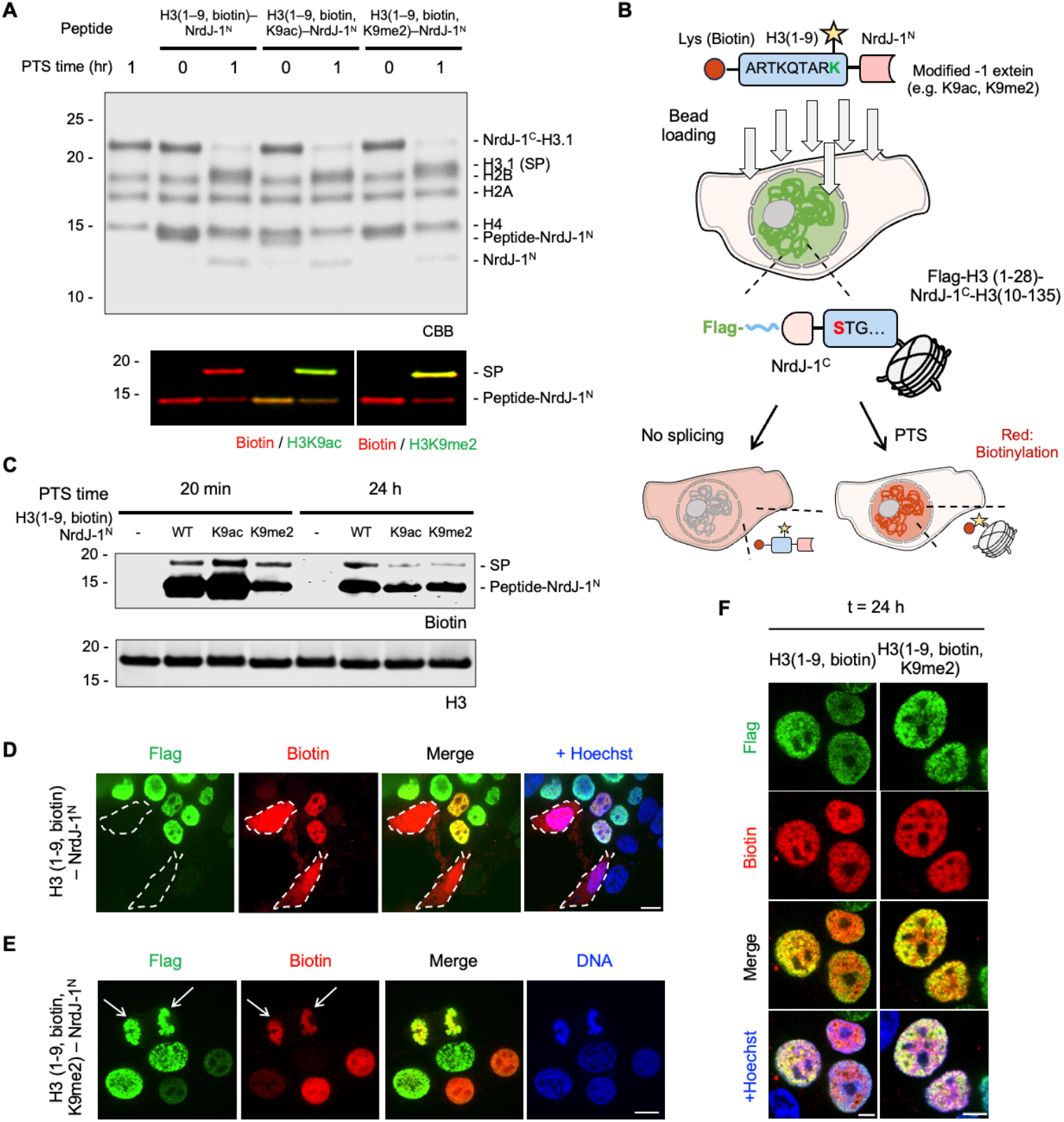
Semisynthesis of modified histone H3 in live cells. (A) *In vitro* PTS reactions on histone octamers containing NrdJ-1^C^–H3(10–135). Purified octamers were reacted with either H3(1–9, biotin)- NrdJ-1^N^, H3(1–9, biotin, K9ac)-NrdJ-1^N^ or H3(1–9, biotin, K9me2)-NrdJ-1^N^. Reaction mixtures were analyzed at indicated timepoint by western blot (bottom) and SDS-PAGE (top). SP, spliced product. (B) Schematic of live-cell PTS via bead loading of modified intein substrates. Cells lacking Flag-H3(1-28)- NrdJ-1^C^-H3.1(10-135) expression display diffuse biotin signal (red) distributed throughout the nucleus and cytosol after bead loading. In contrast, cells expressing Flag–H3(1-28)-NrdJ-1^C^-H3.1(10-135) exhibit nuclear colocalization of Flag (green) and biotin (red) as a result of productive chromatin-associated PTS. (C) Live cell PTS reactions monitored by western blotting. Transfected HEK293T cells were bead loaded with indicated H3(1–9)-NrdJ-1^N^ constructs and then analyzed for biotin content by streptavidin blotting at indicated timepoints. (D-E) Confocal microscopy of HEK293T cells transiently expressing Flag–H3(1-28)-NrdJ-1^C^-H3.1(10-135). Biotinylated H3(1–9)-NrdJ-1^N^ (D) or biotinylated H3(1–9, K9me2)-NrdJ-1^N^ (E) peptides were introduced into the cultured cells by bead loading 24 h post-transfection. Cells were fixed 20 mins post addition of peptides. Immunofluorescence staining was performed with anti-Flag (green) and neutravidin (red). In panel D, two non-transfected cells are outlined with dashed lines. DNA is stained with Hoechst (blue). Arrows indicate a cell in anaphase. Scale bar, 10 μm. (F) Confocal microscopy of HEK293T cells expressing Flag-H3(1-28)-NrdJ-1C-H3.1(10-135). Biotinylated H3(1–9)-NrdJ-1^N^ or biotinylated H3(1–9, K9me2)-NrdJ-1^N^ peptides were introduced into cultured cells by bead loading 24 h post-transfection. Immunostaining was performed as described in panel C but after 24 h. Scale bar, 5 μm.

Encouraged by these results, we delivered modified H3(1–9)-NrdJ-1^N^ conjugates bearing PTMs into cells expressing the complementary NrdJ-1^C^-histone fusion and evaluated splicing after 20 min incubation using the biotin detection handle (Figure 5B). Western blotting revealed successful PTS-mediated semisynthesis had occurred upon delivery of the H3(1-9, biotin)-NrdJ-1^N^, H3(1-9, biotin, K9ac)-NrdJ-1^N^ and H3(1-9, biotin, K9me2)-NrdJ-1^N^ (Figure 5C). Moreover, immunofluorescence revealed robust nuclear signals only in those cells expressing Flag-H3(1-28)-NrdJ-1^C^-H3.1(10-135), both for the biotinylated substrate and for the substrate bearing biotin/H3K9me2 dual modifications, whereas no biotin signal was detected in the negative controls (Figure 5D-E, and Figure S16A-B). Notably, we also detected strong biotin enrichment on mitotic chromatin (e.g. cells undergoing anaphase in Figure 5D and Figure S16B), suggesting that modifications installed on chromatin using PTS might be maintained through cell division. To this point, the biotin signal remained detectable 24 h after splicing

(Figure 5C, 5F and Figure S16C). Overall, these findings establish NrdJ-1-mediated PTS as a practical strategy for modifying endogenous proteins and chromatin in living cells while preserving native primary sequence.

## Conclusions

The discovery of ultrafast split inteins has greatly expanded the scope of PTS in the protein engineering area. Here, we focused on one such split intein, NrdJ-1, focusing on its ability to support protein semisynthesis without the need to introduce facilitating mutations in the target sequence, as often the case when using split inteins^4,22,29^. Our studies show that the exceptional splicing efficiency of NrdJ-1 is largely independent of local extein sequences. This along with its use of a serine at the +1 extein position makes NrdJ-1 well suited to the traceless installation of ncAAs and PTMs into target proteins. As a demonstration of this, we used NrdJ-1 to install various modifications into histone H3 sequence both *in vitro* and in cells. The ability to edit native chromatin without the need to make mutations in the primary sequence opens new avenues for studying chromatin function. In particular, the rapid kinetics of NrdJ-1 PTS should allow for temporally resolved studies of PTM dynamics, such as monitoring the appearance, disappearance, turnover or redistribution of specific modifications over defined time windows. Importantly, split inteins from different families share only structural homology but not primary sequence identity, so that they do not cross-react^19,41^. This inherent orthogonality enables multiplexed applications in which multiple engineered split inteins, including NrdJ-1, can be used in combination without interference. The current studies, by surveying the scope of NrdJ-1 PTS, are expected to accelerate such efforts.

## Supporting information

The Supplementary Information includes Supplementary Figures S1-S16, Supplementary Table S1, Materials and Methods, and additional construct details

## Author Contributions

XY and TWM conceived the project, analyzed the data and wrote the manuscript. Experiments were performed by XY with the help of JS, CK, AD and JZ. TWM supervised the project.

## Data Availability

All data are included in the main figures or supplemental information (SI).References cited in the SI have been listed at the bottom of the main article’s reference list.

## Acknowledgements

We thank member of the Muir lab for valuable discussions. This work was supported by the National Institutes of Health (NIH-GMS grant R01 GM086868) and by funds from the Ludwig Institute for Cancer Research and the Princeton Catalysis Initiative. CK was supported by an EMBO postdoctoral fellowship. JZ is a Damon Runyon Fellow supported by the Damon Runyon Cancer Research Foundation (DRG-2506-23).

