## Supplementary material for "Traceless protein semi-synthesis in cells using the promiscuous ultra-fast split intein NrdJ-1": The Supplementary Information includes Supplementary Figures S1-S16, Supplementary Table S1, Materials and Methods, and additional construct details

**This PDF file includes:**

Supplementary Figures S1-S16 ........ S2

Supplementary Table S1 ........ S24

Materials and Methods ........ S26

DNA constructs and protein sequences ........ S35

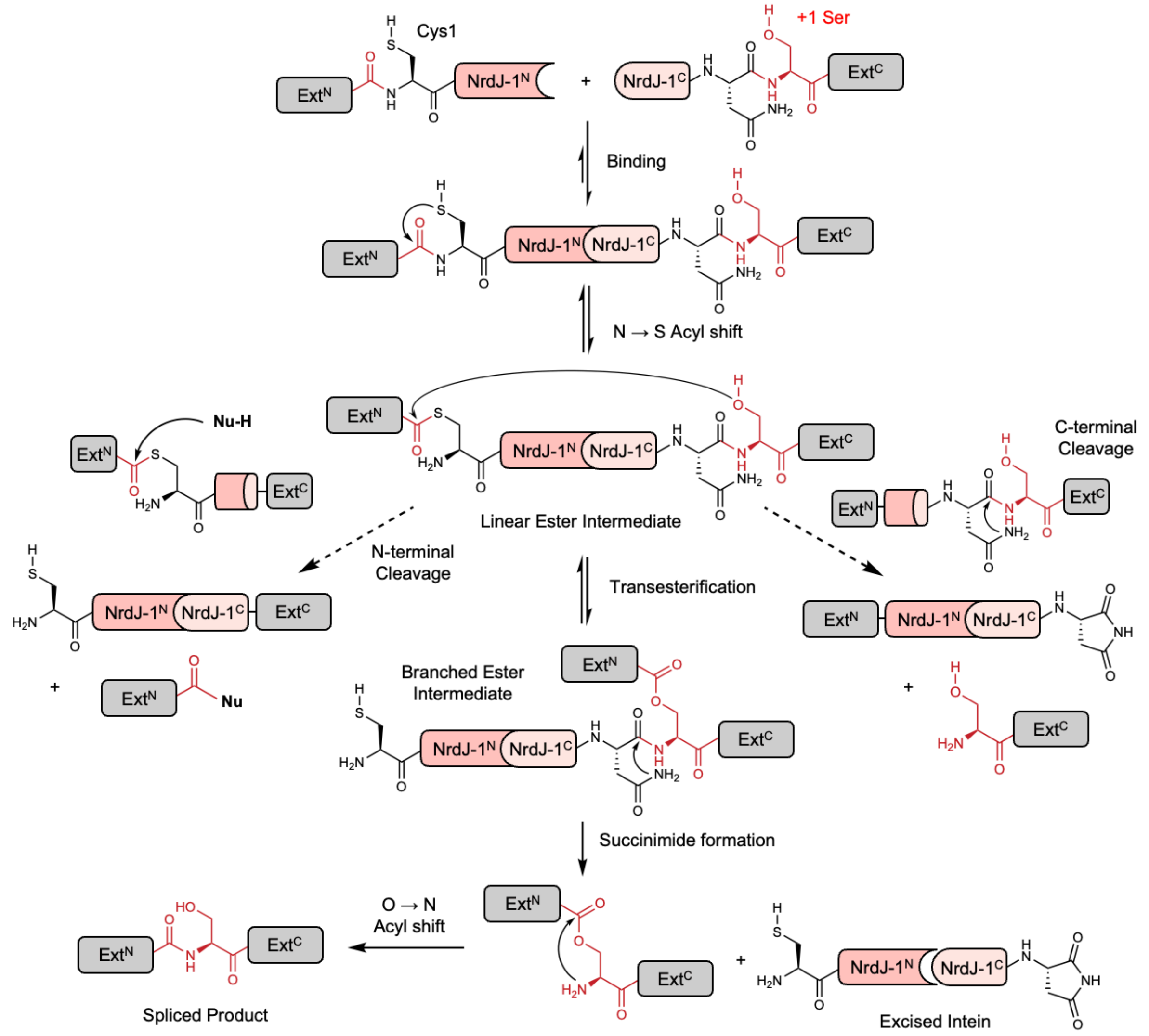


**Supplementary Figure 1. Mechanism of NrdJ-1-mediated protein splicing and extein-dependent side reactions.** Biochemical mechanism of protein splicing and associated N- and C-terminal cleavage pathways with split intein NrdJ-1.

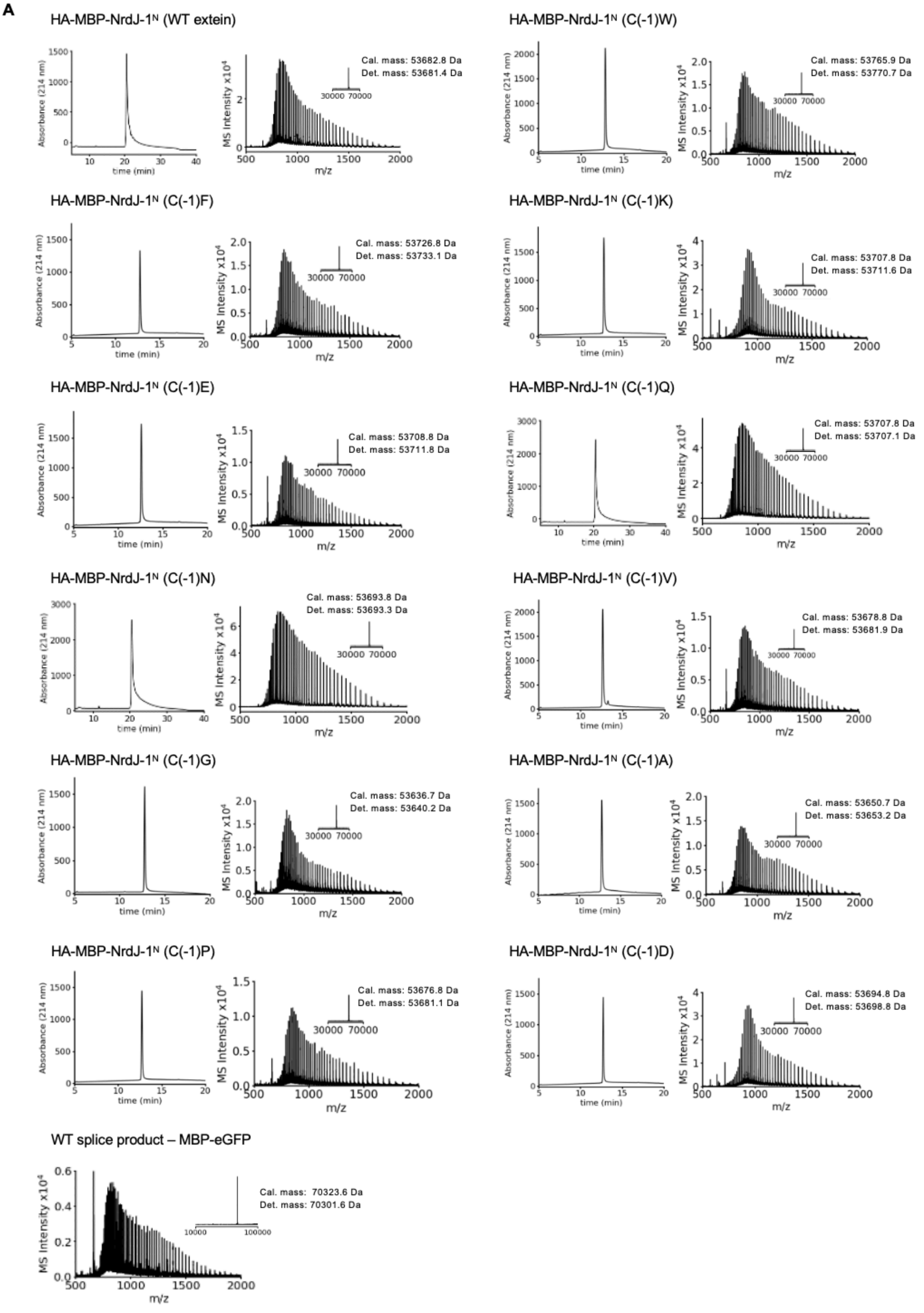
A
HA-MBP-NrdJ-1N (WT extein)
Cal. mass: 53682.8 Da
Det. mass: 53681.4 Da
HA-MBP-NrdJ-1N (C(-1)W)
Cal. mass: 53765.9 Da
Det. mass: 53770.7 Da
HA-MBP-NrdJ-1N (C(-1)F)
Cal. mass: 53726.8 Da
Det. mass: 53733.1 Da
HA-MBP-NrdJ-1N (C(-1)K)
Cal. mass: 53707.8 Da
Det. mass: 53711.6 Da
HA-MBP-NrdJ-1N (C(-1)E)
Cal. mass: 53708.8 Da
Det. mass: 53711.8 Da
HA-MBP-NrdJ-1N (C(-1)Q)
Cal. mass: 53707.8 Da
Det. mass: 53707.1 Da
HA-MBP-NrdJ-1N (C(-1)N)
Cal. mass: 53693.8 Da
Det. mass: 53693.3 Da
HA-MBP-NrdJ-1N (C(-1)V)
Cal. mass: 53678.8 Da
Det. mass: 53681.9 Da
HA-MBP-NrdJ-1N (C(-1)G)
Cal. mass: 53636.7 Da
Det. mass: 53640.2 Da
HA-MBP-NrdJ-1N (C(-1)A)
Cal. mass: 53650.7 Da
Det. mass: 53653.2 Da
HA-MBP-NrdJ-1N (C(-1)P)
Cal. mass: 53676.8 Da
Det. mass: 53681.1 Da
HA-MBP-NrdJ-1N (C(-1)D)
Cal. mass: 53694.8 Da
Det. mass: 53698.8 Da
WT splice product – MBP-eGFP
Cal. mass: 70323.6 Da
Det. mass: 70301.6 Da
Absorbance (214 nm)
time (min)
MS Intensity x10^4
m/z

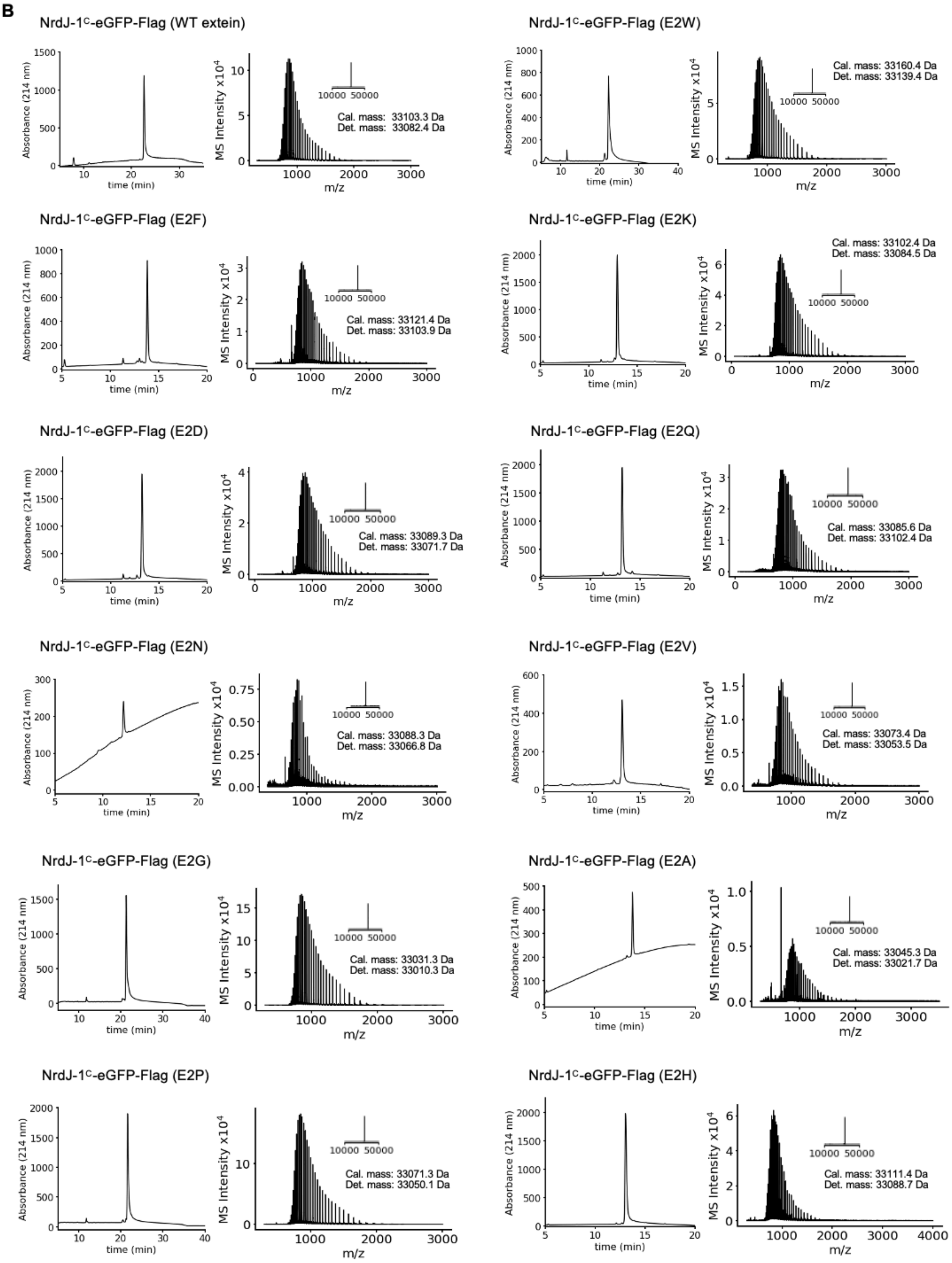
B
NrdJ-1C-eGFP-Flag (WT extein)
Cal. mass: 33103.3 Da
Det. mass: 33082.4 Da
NrdJ-1C-eGFP-Flag (E2W)
Cal. mass: 33160.4 Da
Det. mass: 33139.4 Da
NrdJ-1C-eGFP-Flag (E2F)
Cal. mass: 33121.4 Da
Det. mass: 33103.9 Da
NrdJ-1C-eGFP-Flag (E2K)
Cal. mass: 33102.4 Da
Det. mass: 33084.5 Da
NrdJ-1C-eGFP-Flag (E2D)
Cal. mass: 33089.3 Da
Det. mass: 33071.7 Da
NrdJ-1C-eGFP-Flag (E2Q)
Cal. mass: 33085.6 Da
Det. mass: 33102.4 Da
NrdJ-1C-eGFP-Flag (E2N)
Cal. mass: 33088.3 Da
Det. mass: 33066.8 Da
NrdJ-1C-eGFP-Flag (E2V)
Cal. mass: 33073.4 Da
Det. mass: 33053.5 Da
NrdJ-1C-eGFP-Flag (E2G)
Cal. mass: 33031.3 Da
Det. mass: 33010.3 Da
NrdJ-1C-eGFP-Flag (E2A)
Cal. mass: 33045.3 Da
Det. mass: 33021.7 Da
NrdJ-1C-eGFP-Flag (E2P)
Cal. mass: 33071.3 Da
Det. mass: 33050.1 Da
NrdJ-1C-eGFP-Flag (E2H)
Cal. mass: 33111.4 Da
Det. mass: 33088.7 Da
Absorbance (214 nm)
time (min)
MS Intensity x10^4
m/z
10000 50000

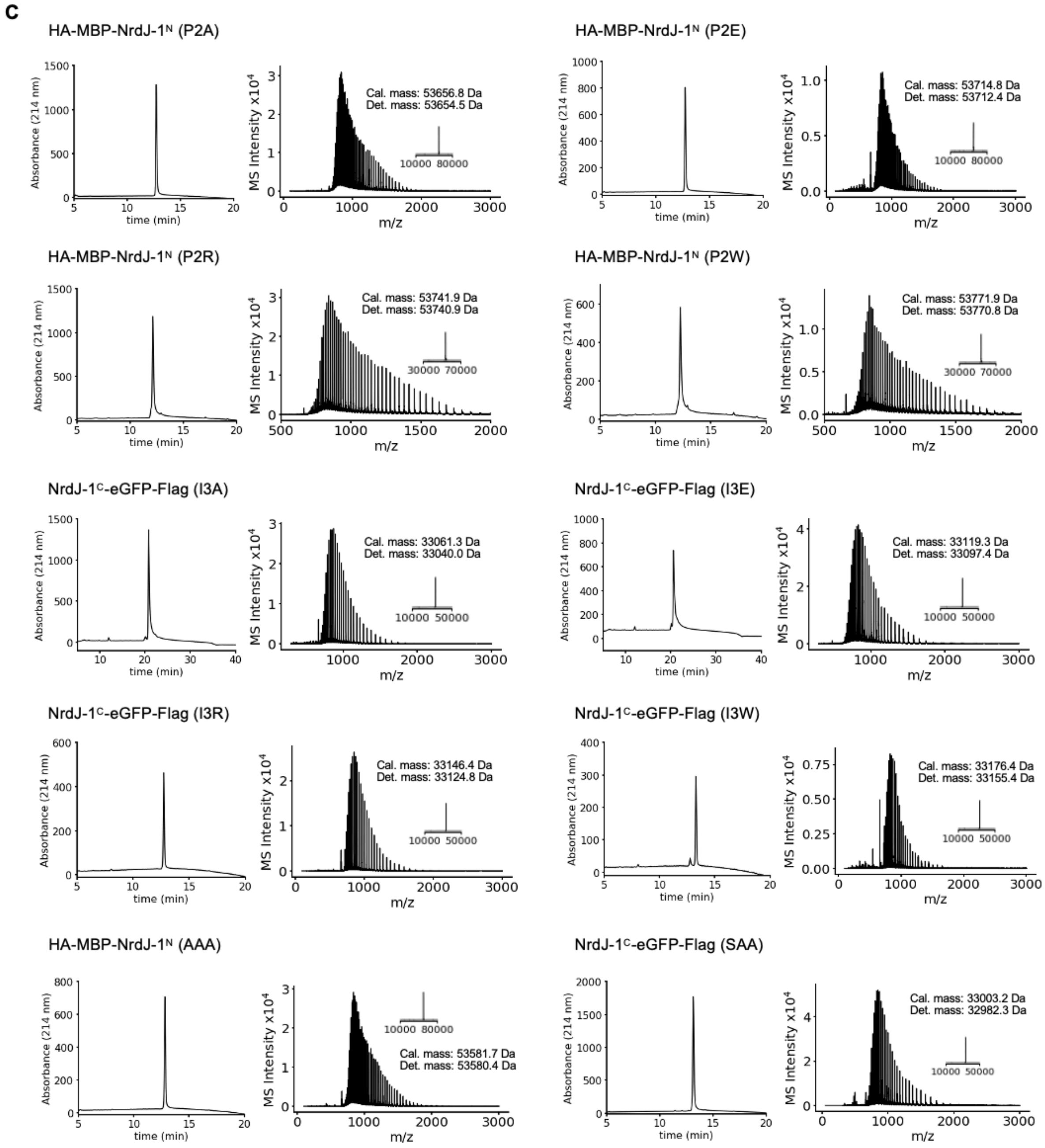


**Supplementary Figure 2. Protein characterization of NrdJ-1 extein constructs**. (A-C) Purified proteins and the wildtype (WT) spliced product were analyzed by RP-HPLC (left) using a 0–100% B gradient over 30 or 40 min, and by ESI-TOF MS (right). The expected and observed molecular masses are indicated for (A) WT int$^{N}$, −1 extein constructs and WT spliced product. (B) WT int$^{C}$ and +2 extein constructs. (C) −2/+3 extein constructs. The ~20.8 Da mass shift observed for the NrdJ-1$^{C}$–eGFP constructs is attributed to eGFP chromophore maturation.

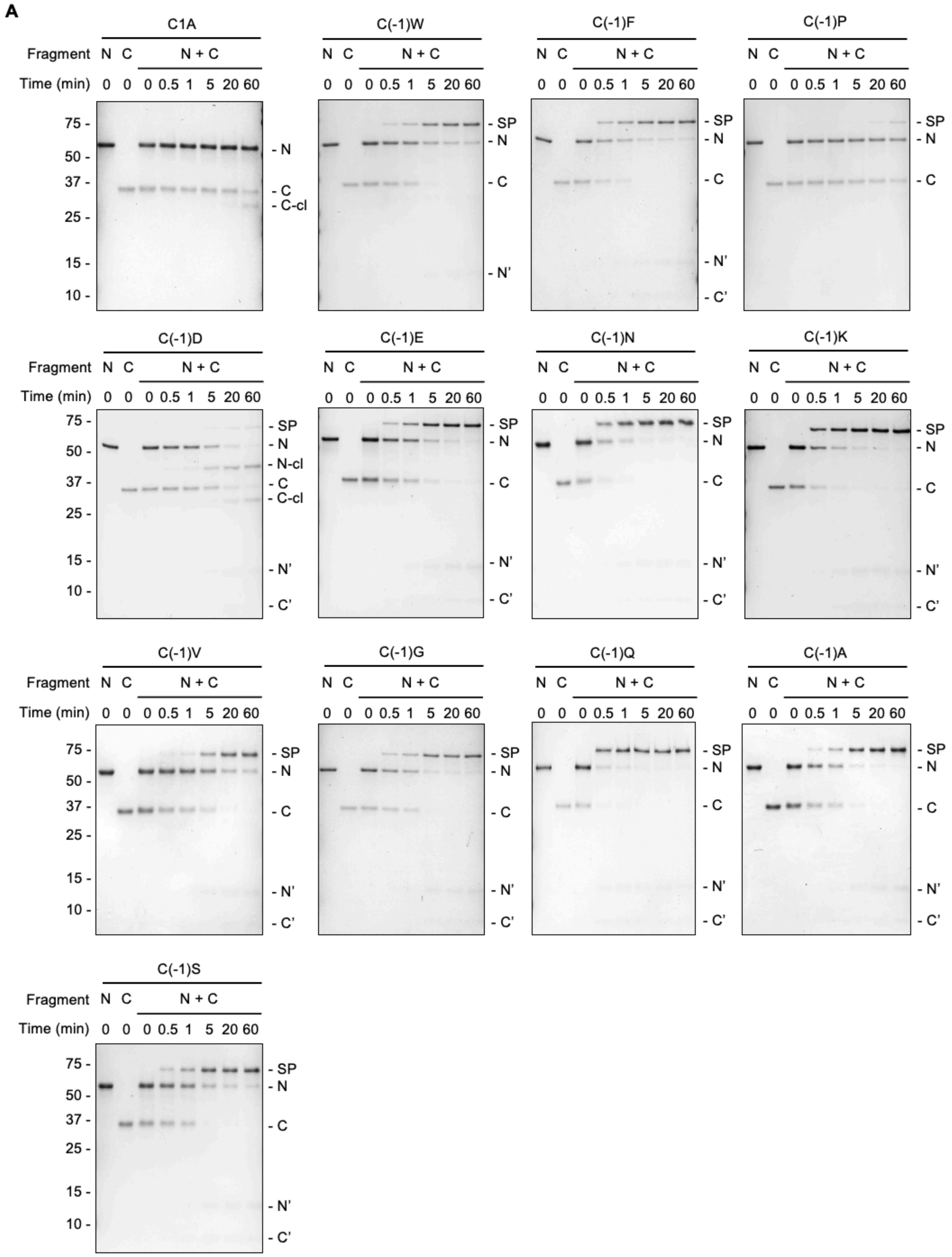
A
C1A
C(-1)W
C(-1)F
C(-1)P
C(-1)D
C(-1)E
C(-1)N
C(-1)K
C(-1)V
C(-1)G
C(-1)Q
C(-1)A
C(-1)S
Fragment N C N + C
Time (min) 0 0 0 0.5 1 5 20 60
75 -
50 -
37 -
25 -
15 -
10 -
- SP
- N
- N-cl
- C
- C-cl
- N'
- C'

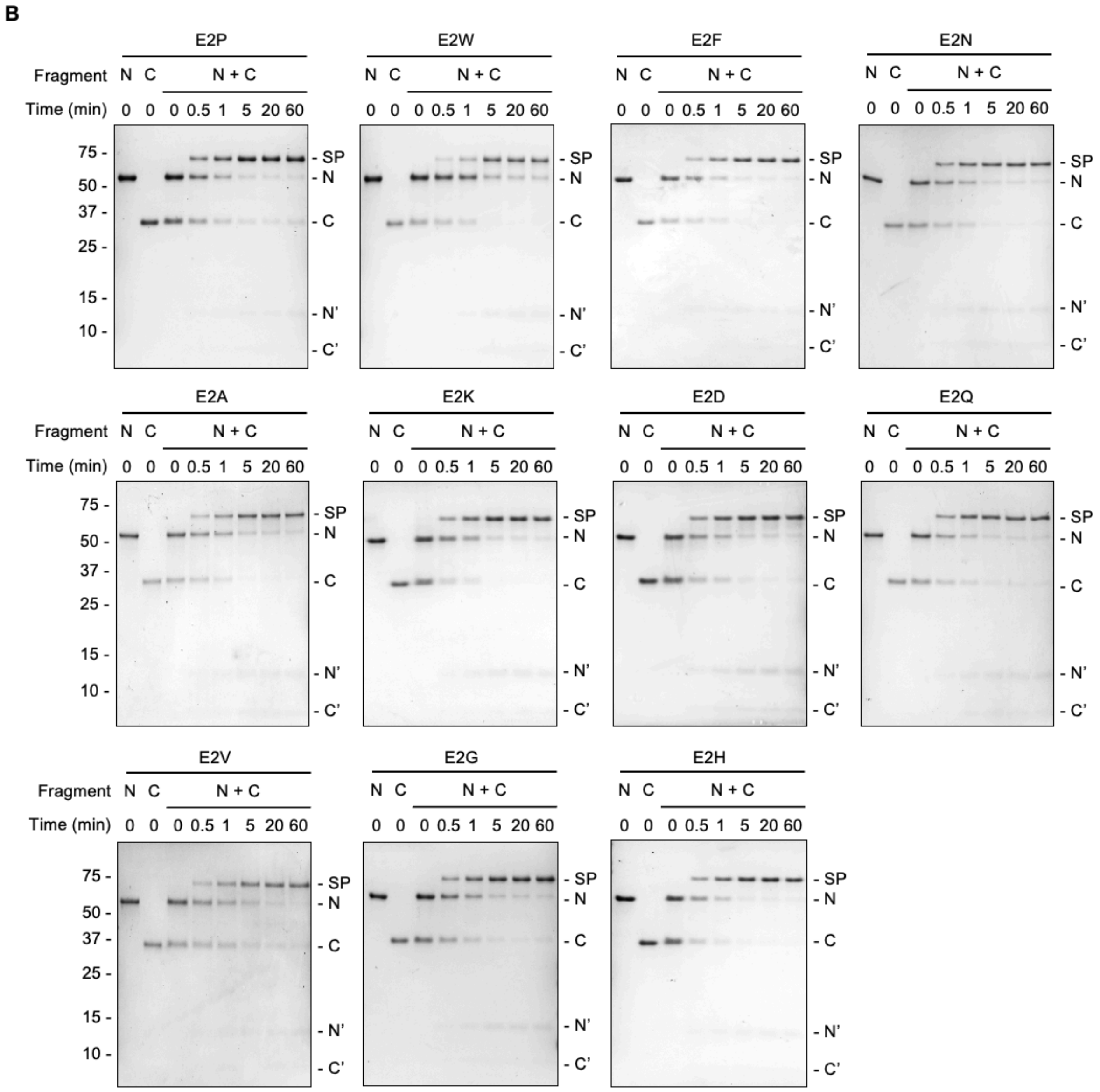
B
E2P
E2W
E2F
E2N
E2A
E2K
E2D
E2Q
E2V
E2G
E2H
Fragment
N
C
N + C
Time (min)
0 0 0 0.5 1 5 20 60
75 -
50 -
37 -
25 -
15 -
10 -
- SP
- N
- C
- N'
- C'

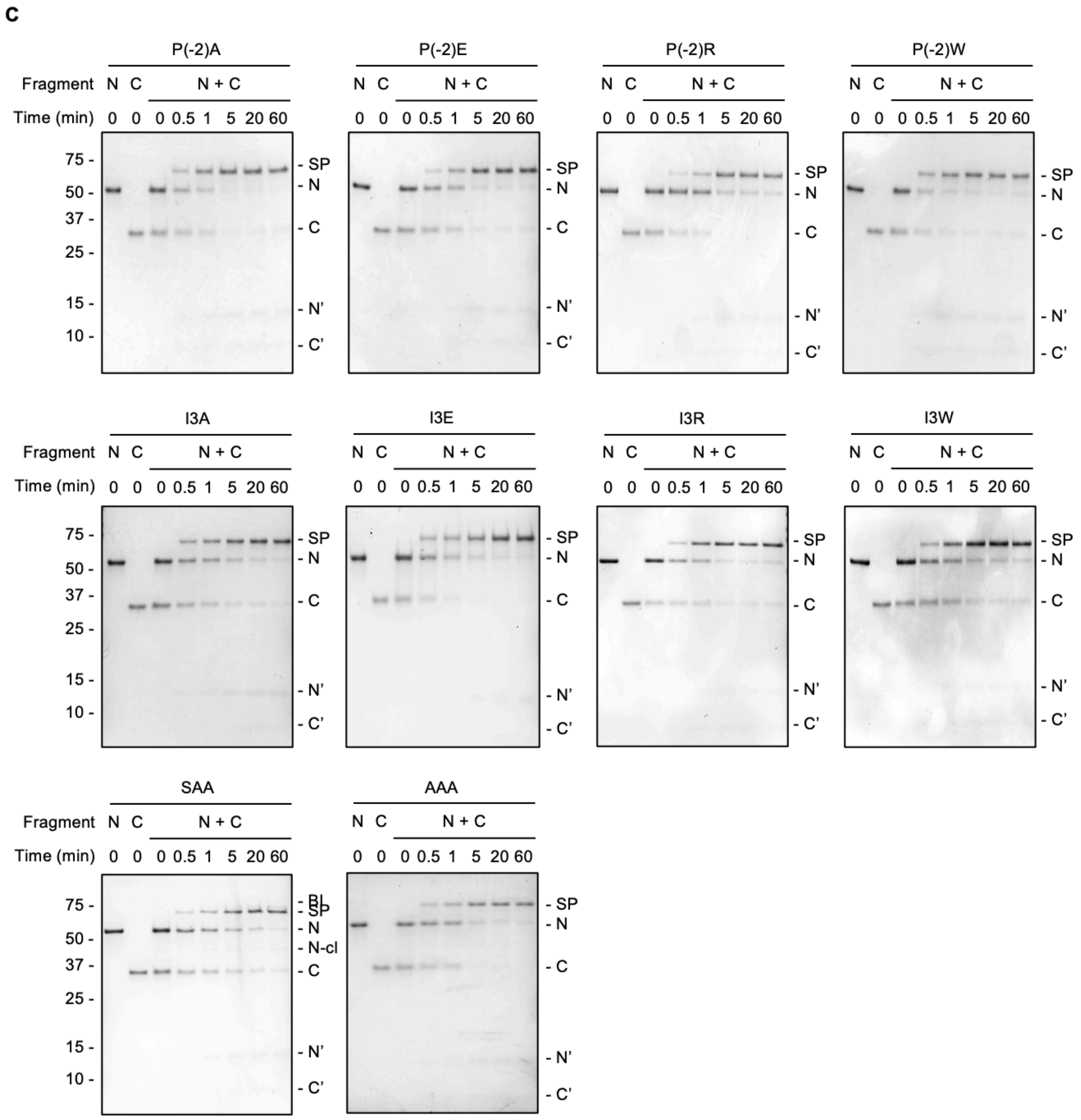


**Supplementary Figure 3. *In vitro* PTS characterization of extein dependence of NrdJ-1.** (A-C) Representative gels for *in vitro* PTS splicing bearing different extein residues in figure 2, quenched at indicated timepoints. Splicing reactions are visualized by SDS-PAGE followed by Coomassie staining. SP: spliced product; N: MBP–$X_{-2}X_{-1}$–NrdJ-1$^{N}$; N-cl: N-terminal cleavage; C: NrdJ-1$^{C}$– $X_1X_2X_3$–eGFP; C-cl: C-terminal; N’: N intein after splicing; C’: C intein after splicing. Panel A corresponds to varying -1 extein residue, and C1A negative control; Panel B corresponds to varying +2 extein residues and panel C corresponds to varying other extein residues as indicated.

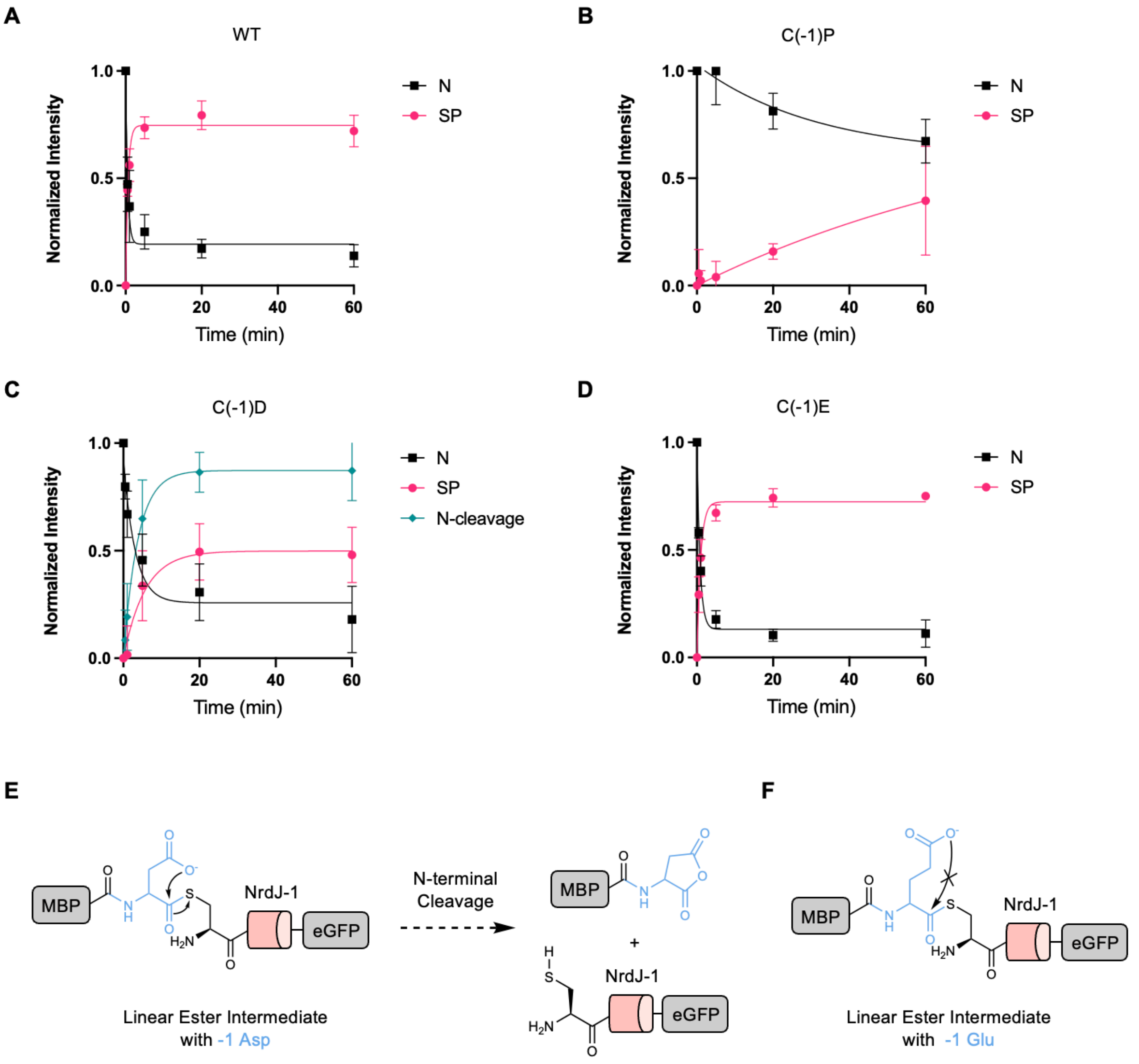


**Supplementary Figure 4. Kinetic curves for reaction progress analysis and proposed mechanism for N-terminal cleavage in -1 Asp construct.** (A–D) Reaction progress curves showing the normalized abundance of the starting material (N; MBP-X$_{-2}$/X$_{-1}$-NrdJ-1$^{N}$) and splice products (SP) over time. N-terminal cleavage was monitored for constructs containing -1 Asp extein. (A) WT (-1 Cys), (B) -1 Pro, (C) -1 Asp, and (D) -1 Glu. Error bars indicate SD (n = 3). (E-F) Proposed mechanisms for N-terminal cleavage side-product formation with -1 Asp extein. -1 Asp facilitates the formation of a 5-membered cyclic anhydride, causing internal cleavage of the N-extein. Similar side reaction was not observed with -1 Glu, which would form a 6-membered ring.

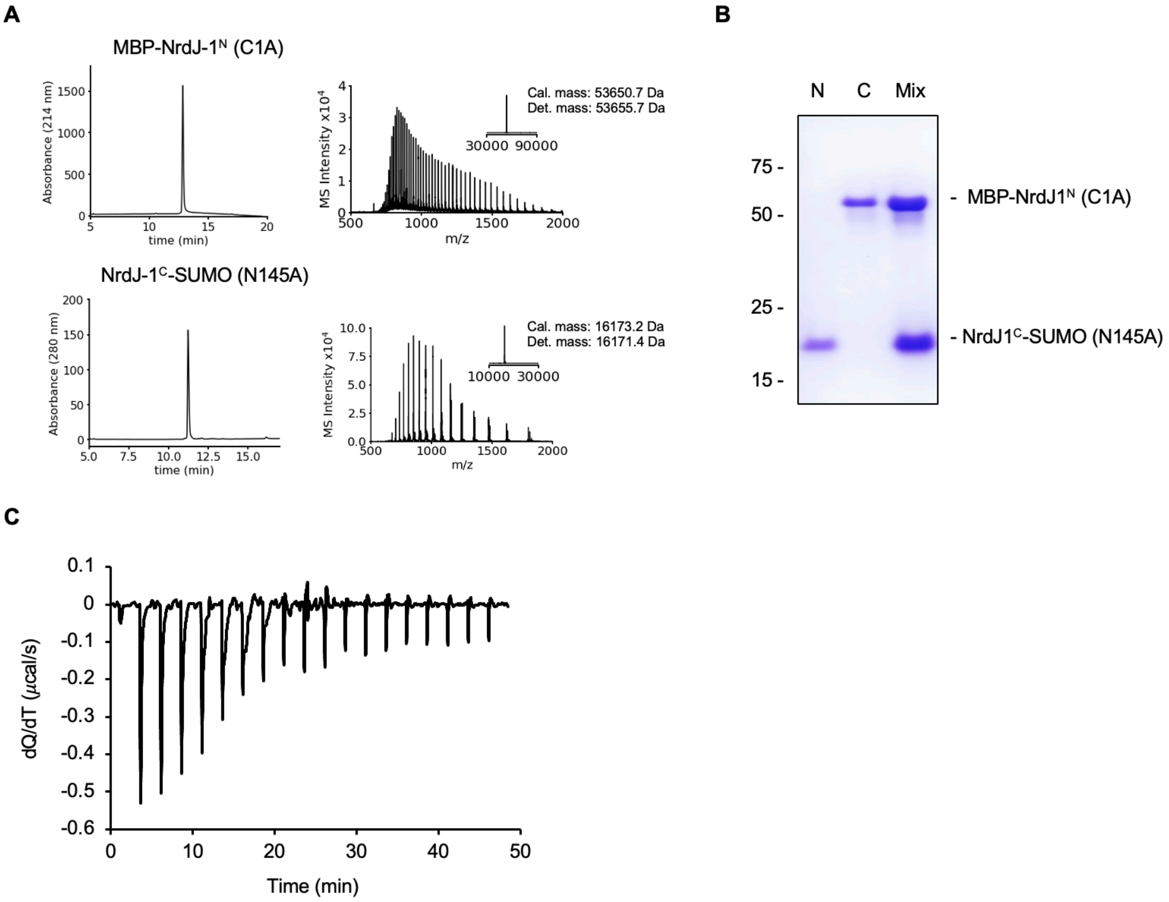


**Supplementary Figure 5. Isothermal calorimetry for the NrdJ-1 split intein.** (A) RP-HPLC analysis (0-100% B gradient over 30 min) and ESI-TOF MS of NrdJ-1 constructs used in the ITC study. (B) SDS-PAGE analysis of the individual and mixed intein fragments. The two inactivated fragments do not undergo splicing upon mixing. (C) Titration curve of the reaction corresponding to Figure 3A, showing the heat change (dQ/dT) over time during titration of NrdJ-1$^{C}$-SUMO (N145A) into a solution containing MBP-NrdJ-1$^{N}$(C1A).

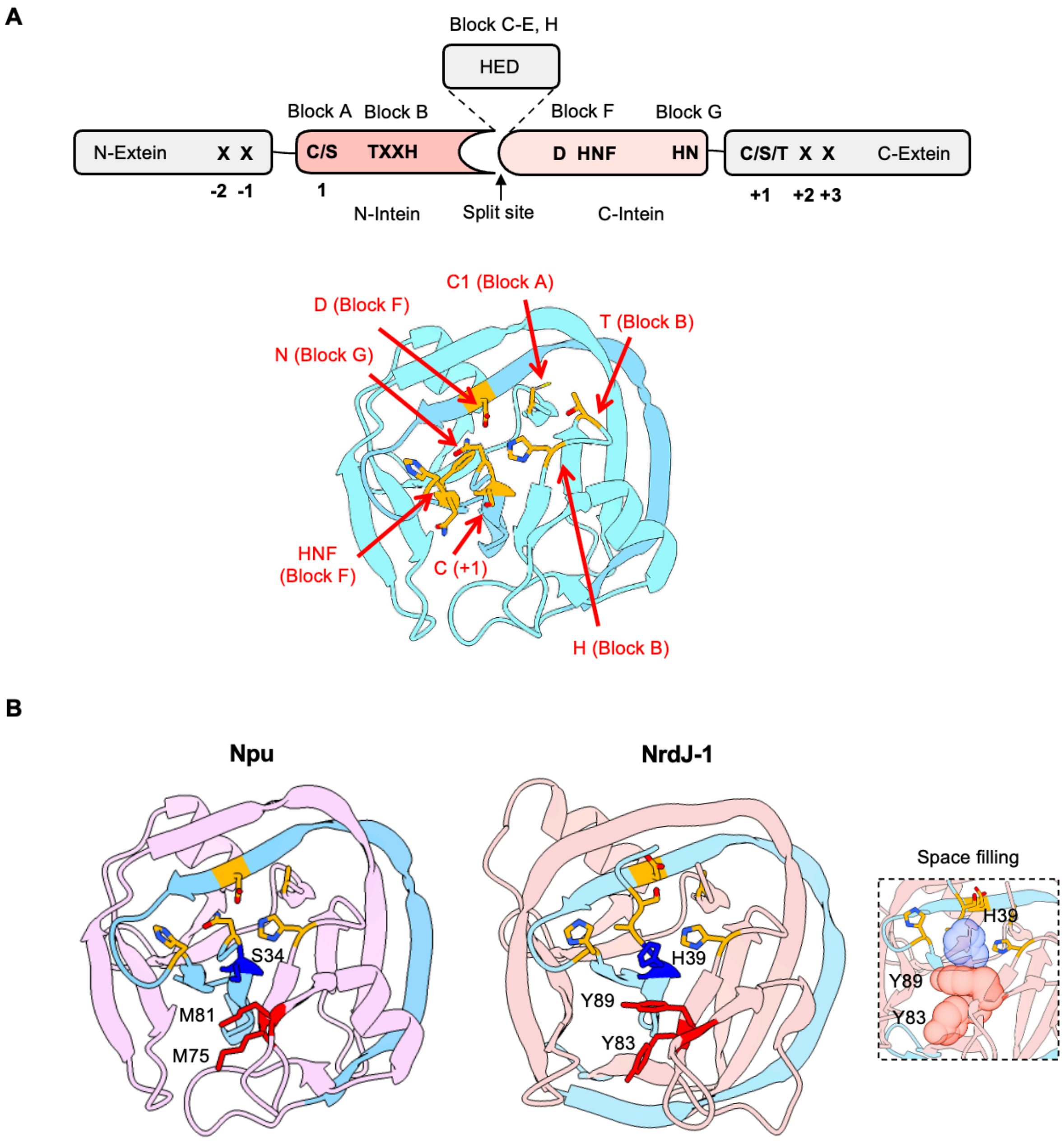


**Supplementary Figure 6. Conserved sequence blocks and accelerator residues in split intein structures.** (A) Top: Inteins have conserved sequence motifs. The homing endonuclease domain (HED) is only present in some contiguous inteins and are absent in split inteins. The first residue of the intein is designated as position 1. The extein junctions flanking the intein (positions –2, –1, +1, +2, +3) can influence splicing efficiency. Bottom: the X-ray protein crystal structure of Npu with conserved catalytic residues in the HINT fold highlighted in orange and labelled. (B) Comparison of the accelerator residues of NrdJ-1 and Npu split inteins. The regions corresponding to the N- and C-intein fragments in the split version are colored pink and blue, respectively. The accelerator residues residing in the N-terminal and the C-terminal split intein fragments are colored red and blue, respectively. Space filling representations of the accelerator residues in NrdJ-1 are displayed and modeled using UCSF Chimera. PDB: 8UBS (NrdJ-1); 4KL5 (Npu).

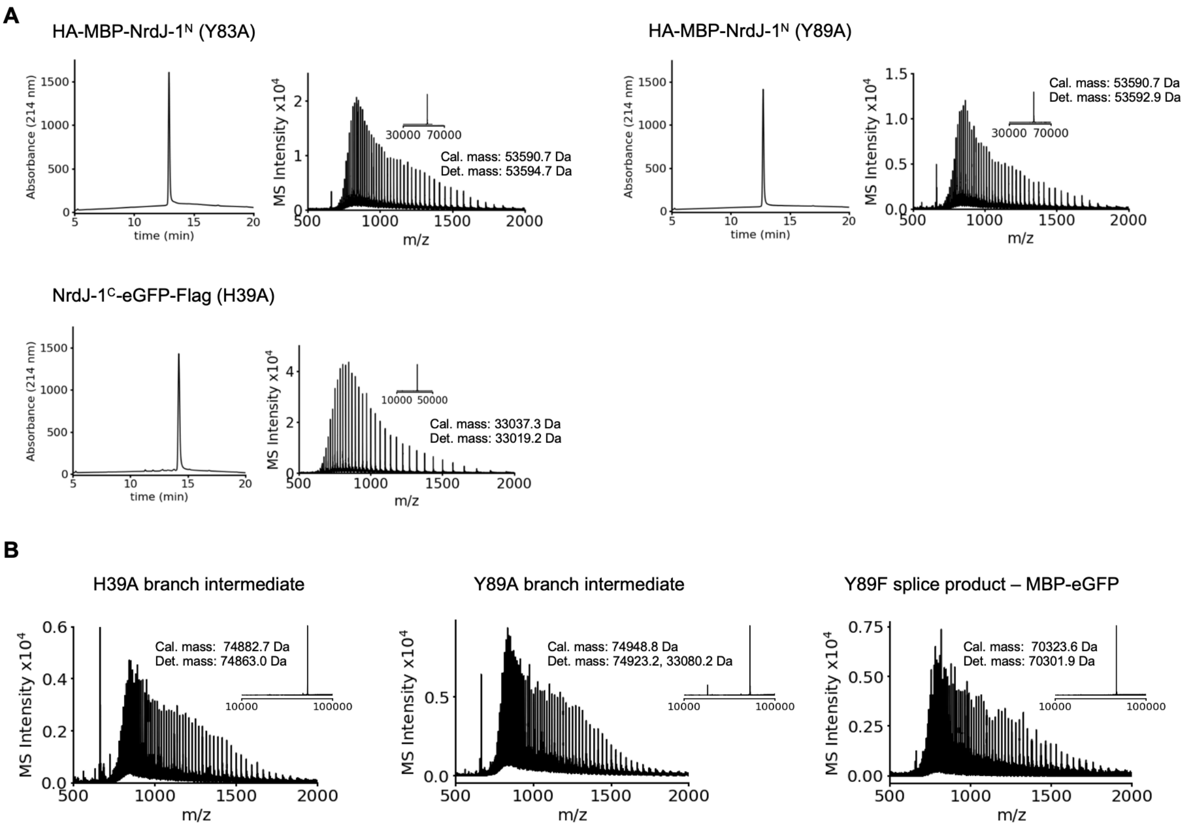


**Supplementary Figure 7. Protein characterization of NrdJ-1 accelerator alanine mutations and their corresponding splice products**. (A) The purified accelerator mutant proteins were analyzed by RP-HPLC (left) using a 0–100% B gradient over 30 min, and by ESI-TOF MS (right). (B) The splicing reactions were analyzed by RP-HPLC using a 0–100% B gradient over 30 min, and the indicated spliced products (SP) or branched intermediate (BI) were characterized by ESI-TOF MS. The expected and observed molecular masses are indicated. Minor additional species for Y89A at 33080.2, correspond to NrdJ-$1^C$-eGFP (see Figure S8E). The mass shift observed for the NrdJ-$1^C$–eGFP construct is attributed to eGFP chromophore maturation.

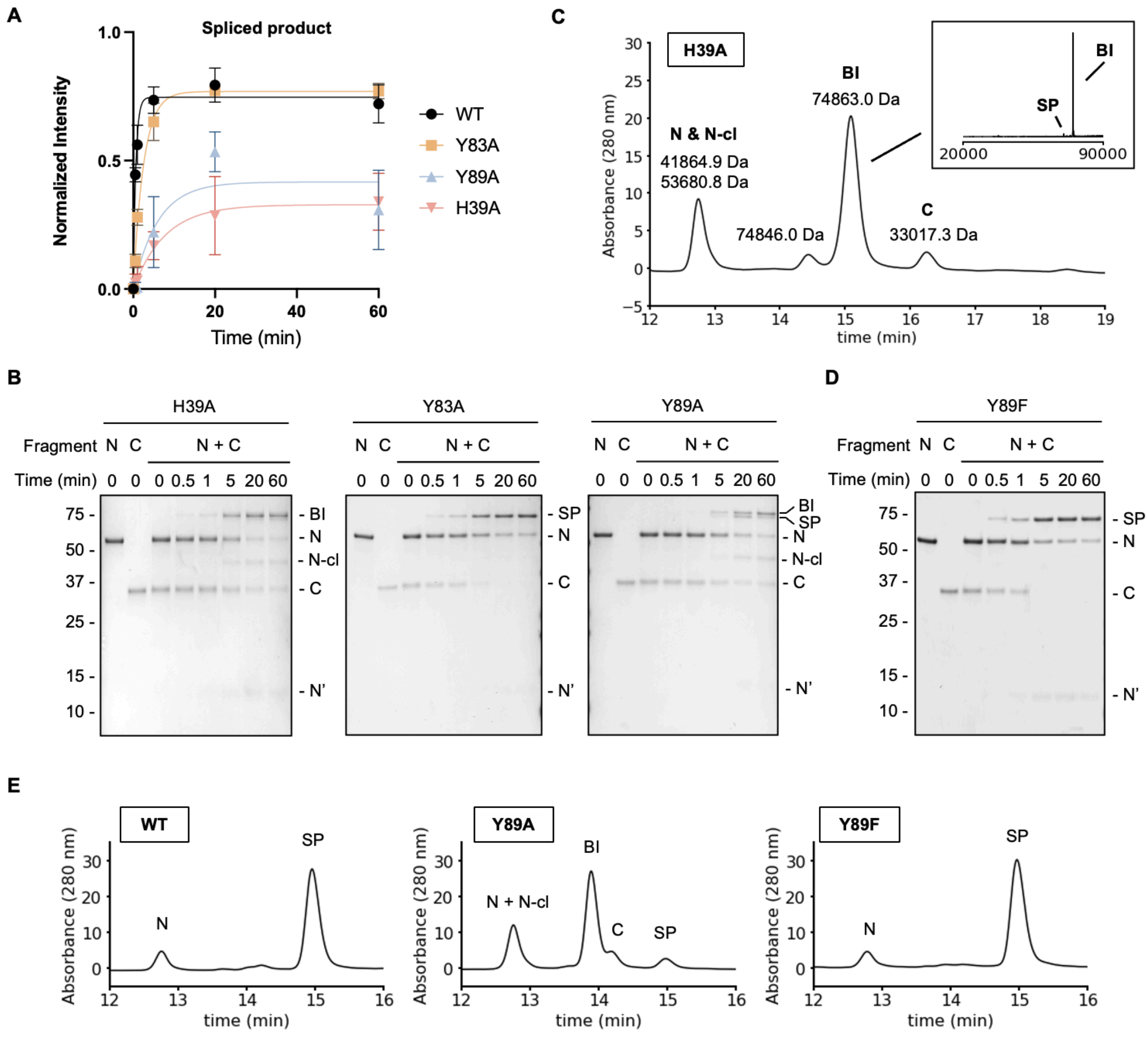


**Supplementary Figure 8. Accelerator residues facilitate succinimide formation.** (A) Reaction progress curves showing the normalized formation of spliced products (SP) over time for accelerator residue mutants. Error bars represent SD (n = 3). (B) Representative SDS-PAGE gels for *in vitro* PTS splicing bearing the different accelerator mutations used in figure 3D, quenched at indicated timepoints. Splicing reactions are visualized by SDS-PAGE followed by Coomassie staining. SP: spliced product; BI: branched intermediate; N: MBP-NrdJ-1N; N-cl: N-terminal cleavage; C: NrdJ-$1^{C}$-eGFP; C-cl: C-terminal cleavage; N': N intein after splicing; C': C intein after splicing. (C) RP-HPLC chromatogram of splice products formed by NrdJ-$1^{C}$-eGFP H39A mutant reacting with wildtype MBP-NrdJ-$1^{N}$ after 30 mins reaction. Identity of peaks was determined by ESI-MS. Inset: zoom-in view of the reconstructed mass spectrum showing majority formation of branched intermediate and minimal formation of splice product. (D) Representative gel for *in vitro* PTS splicing for Y89F mutants. Reaction was carried out as previously described. (E) RP-HPLC analysis of splicing reactions using MBP-NrdJ-$1^{N}$ WT, Y89A, and Y89F reacting with wildtype NrdJ-$1^{C}$-eGFP after 30 mins reaction. Identity of peaks was determined by ESI-MS.

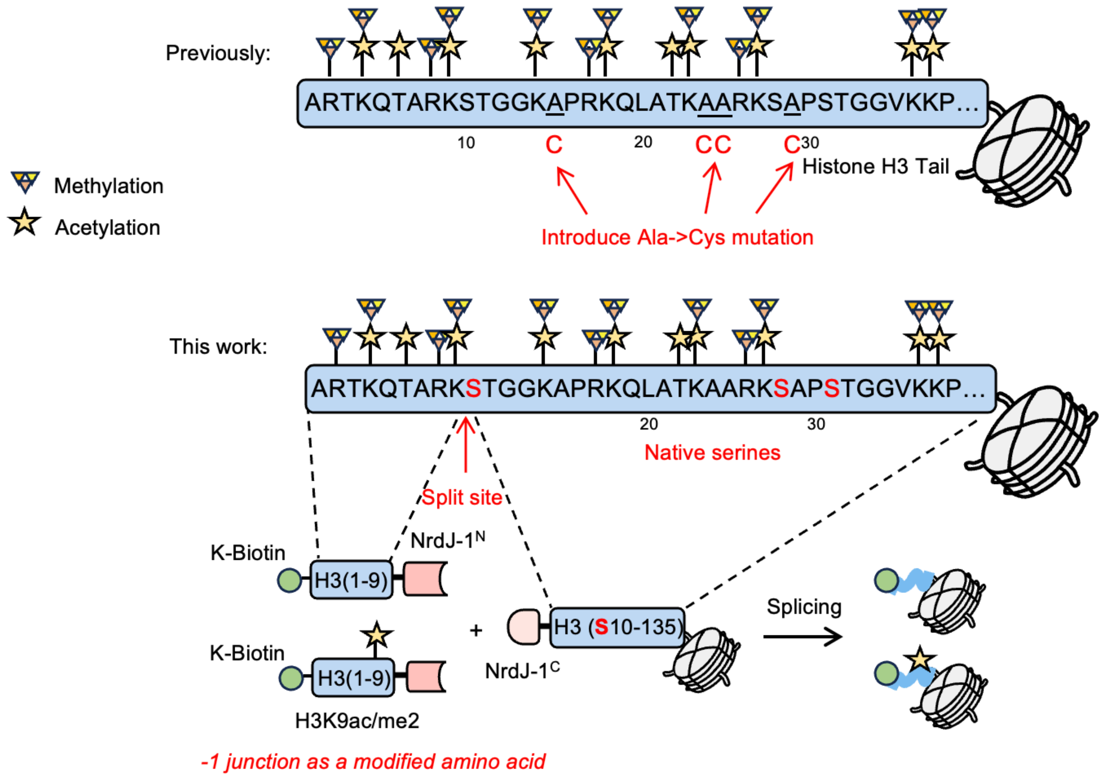


**Supplementary Figure 9. Traceless installation of defined PTMs on the histone H3 N-terminal tail using the serine-based split intein NrdJ-1.** The histone H3 N-terminal tails are heavily modified by various post-translational modifications (PTMs); representative acetylation (stars) and methylation (triangles) are shown. Top: previous approach introduced non-native cysteines (red "C") at selected alanine positions for ligation with cysteine-based inteins, creating point mutations that can be removed by desulfurization; however, desulfurization is incompatible with certain PTMs. Bottom: In this work, we employ the serine-based split intein NrdJ-1 to perform protein trans-splicing (PTS) at native serine residues in H3 (red "S"; S10, S28, S31). A short, PTM-bearing H3(1–9) peptide fused to NrdJ-1N is spliced to a recombinant H3(S10–135) fragment carrying to NrdJ-1C, regenerating full-length H3 within nucleosomes without sequence scars. This strategy enables traceless installation of multiple, user-defined PTMs on the H3 N-terminus.

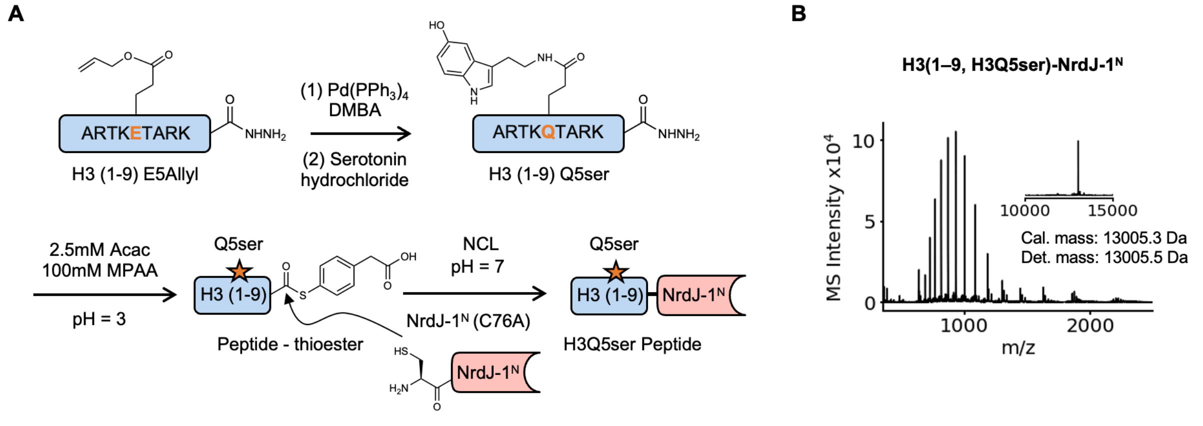


**Supplementary Figure 10. Synthesis of H3(1–9) peptides bearing H3Q5ser ligated to NrdJ-1$^{N}$.** (A) The indicated modified H3(1–9) peptides was generated by SPPS. An E5-allyl precursor peptide was first assembled and then converted to Q5ser. Peptide hydrazide was activated to a C-terminal thioesters and then coupled by NCL to NrdJ-1$^{N}$ (C76A), yielding H3(1–9, Q5ser)–NrdJ-1$^{N}$ ligation products used for subsequent PTS. (B) ESI-TOF MS characterization of H3(1–9, Q5ser)–NrdJ-1$^{N}$(C76A). Inset show deconvoluted masses; calculated and detected masses as indicated.

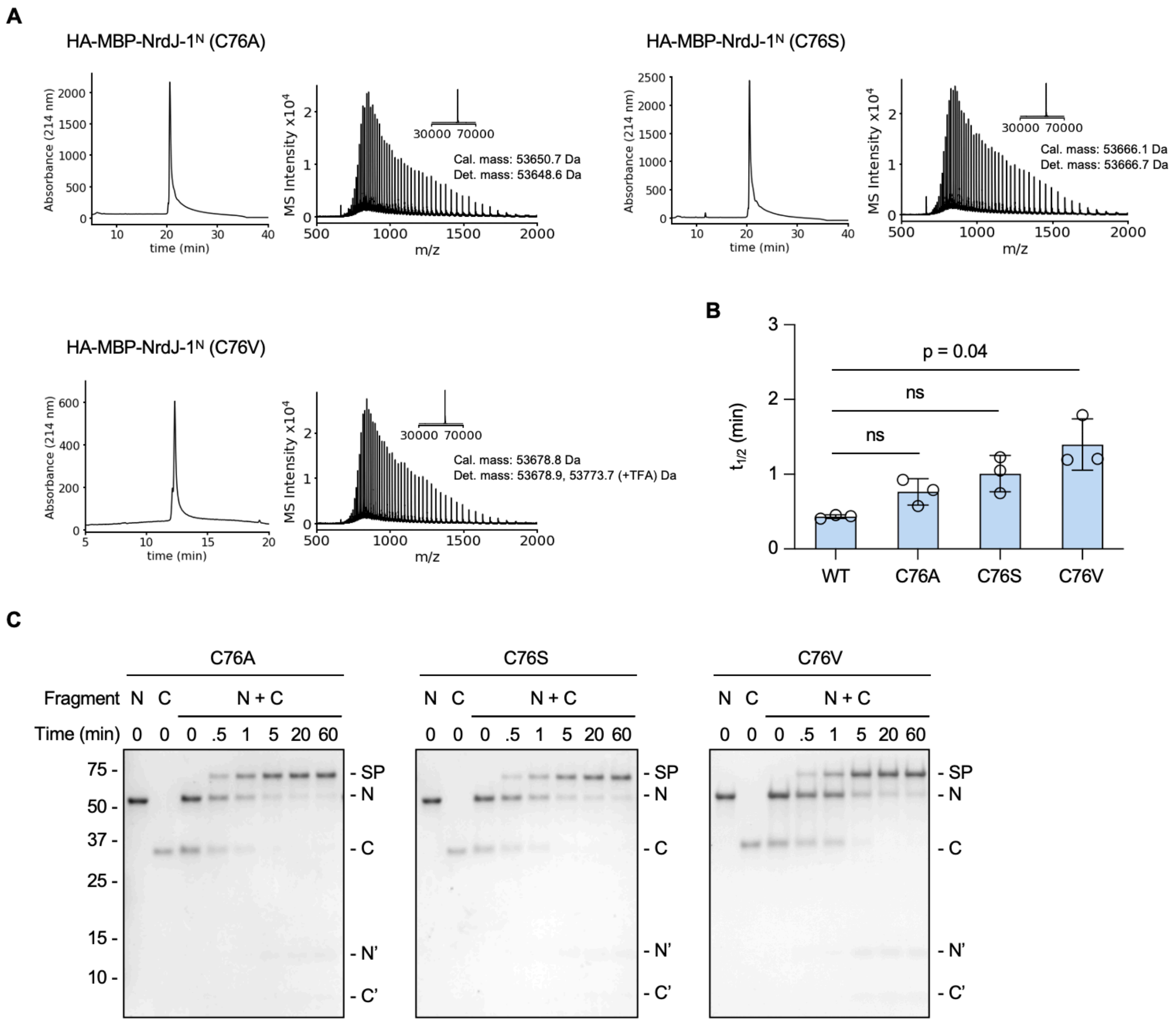


**Supplementary Figure 11. Characterization and splicing of NrdJ-1 Cys76 mutants** (A) The Cys76 mutant proteins purified through size exclusion chromatography were analyzed by RP-HPLC (left) using a 0–100% B gradient over 30 min or 40 min, and by ESI-TOF MS (right). The expected and observed molecular masses are indicated. (B) *In vitro* PTS splicing half-lives for wildtype NrdJ-1 and C76 mutants (C76A, C76S, and C76V). *In vitro* PTS splicing was performed with 1 μM MBP-NrdJ-1$^{N}$, 1 μM NrdJ-1$^{C}$-eGFP in splicing buffer (pH 7.2) and quenched at indicated timepoints. Reaction progress was monitored by SDS-PAGE followed by densitometry analysis of product bands. Error bars indicate standard deviation (SD), n = 3. (C) Representative gels for *in vitro* PTS splicing bearing different C76 mutations. Splicing reactions are visualized by SDS-PAGE followed by Coomassie staining. SP: spliced product; N: MBP-NrdJ-1$^{N}$; C: NrdJ-1$^{C}$-eGFP; N': N intein after splicing; C': C intein after splicing.

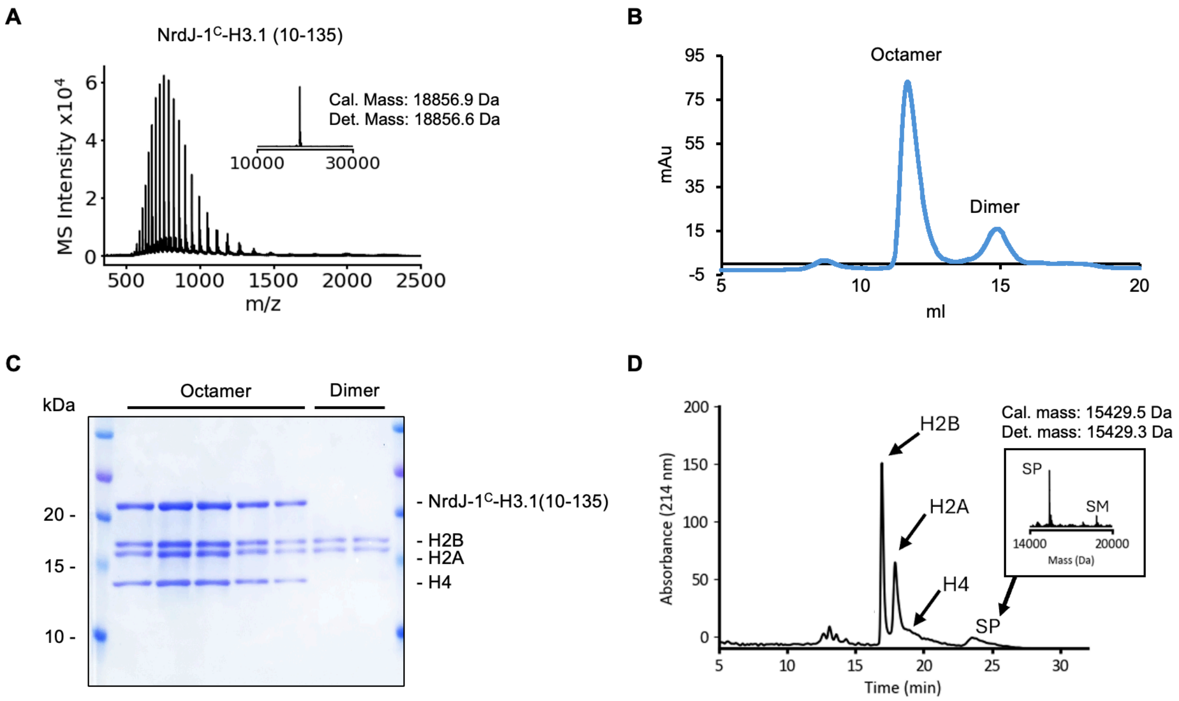


**Supplementary Figure 12. Octamer formation and splicing with NrdJ-1$^{C}$-H3.1 (10-135).** (A) ESI-TOF mass spectrum of purified NrdJ-1$^{C}$-H3.1 (10-135), with calculated and observed masses indicated. Inset shows the deconvoluted mass. (B) Size-exclusion chromatography of the refolded histone mixture. The major octamer peak corresponds to the histone octamer and the minor peak an H2A–H2B dimer. (C) Coomassie-stained SDS-PAGE of SEC fractions for validation of the peak composition resolved by FPLC in panel B. Samples are resolved by a 16% Tris-HCl gel with Coomassie staining. (D) RP-HPLC analysis and the corresponding deconvoluted mass spectra of H3Q5ser octamer splicing reactions. SP, spliced product (H3Q5Ser histone H3); SM, starting material (NrdJ-1$^{C}$–H3.1 [10–135]).

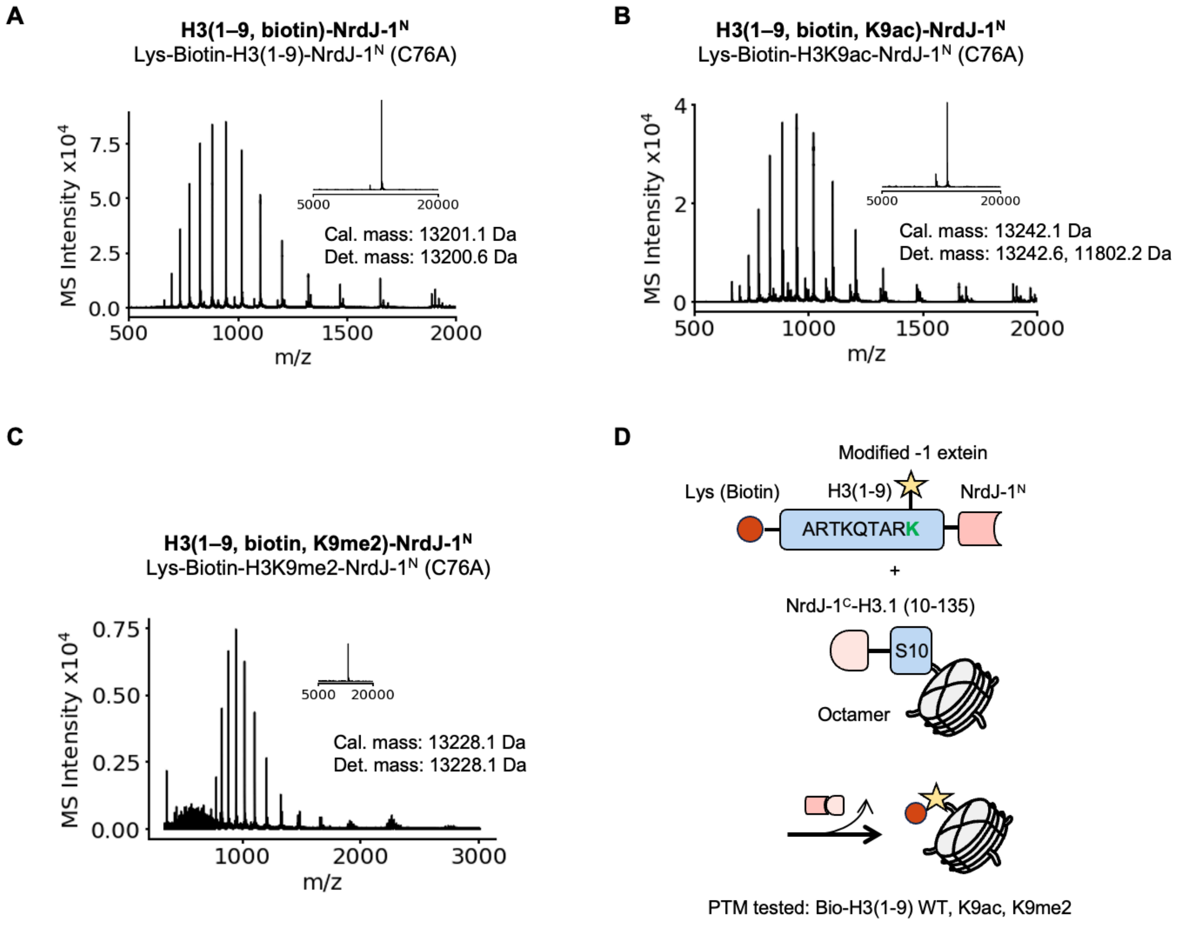


**Supplementary Figure 13. Characterization of PTM-bearing H3(1-9)–NrdJ-1$^{N}$ products.** (A-C) ESI-TOF spectra of the synthetic peptides ligated (via NCL) to NrdJ-1$^{N}$ (C76A). Insets show deconvoluted masses. NCL products were: (A) H3(1-9)-NrdJ-1$^{N}$ (C76A) with a biotinylated lysine at N-terminus; (B) H3(1-9)-NrdJ-1$^{N}$ (C76A) with a biotinylated lysine and H3K9ac (minor additional species at 11,802.2, correspond to NrdJ-1$^{N}$-C76A) at N-terminus; and (C) H3(1-9)-NrdJ-1$^{N}$ (C76A) with a biotinylated lysine and H3K9me2 at N-terminus. (D) Schematic of the *in vitro* splicing on the histone H3 N-terminus tail. Splicing ligates chemically modified H3(1–9) peptides to H3(10–135) within the octamer at native Ser10 and releases the associated split intein fragment. Modification at H3K9 places a PTM-containing lysine directly in front of Ser10, illustrating the extein tolerance of NrdJ-1.

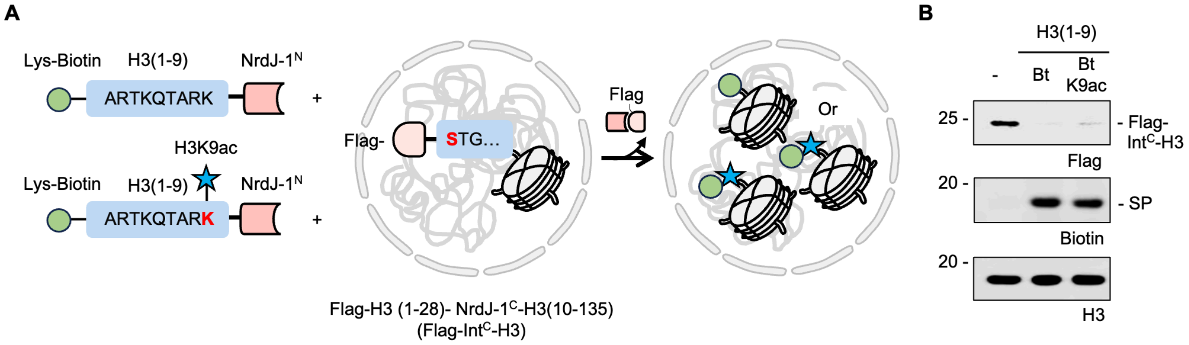


**Supplementary Figure 14. Editing of the H3 N-terminus tail by NrdJ-1 in isolated cell nuclei.** (A) Schematic of the experimental design. Chemically modified substrates H3(1–9, biotin)–NrdJ-1$^{N}$ or H3(1–9, biotin, K9ac)–NrdJ-1$^{N}$ were incubated with isolated HEK293T nuclei transiently expressing Flag-H3(1–28)-NrdJ-1$^{C}$-H3(10–135). Protein *trans*-splicing excises the N-terminal Flag-H3(1–28)-NrdJ-1$^{C}$ and ligates modified H3(1–9) to H3(10–135), installing the PTMs on chromatin; (B) Western blot analysis of nuclear extracts after 1 h incubation at 37 °C. Robust PTS is observed with the disappearance of starting materials (indicated by Flag signal) and the formation of the spliced products (biotinylated signal). Histone H3 serves as loading control.

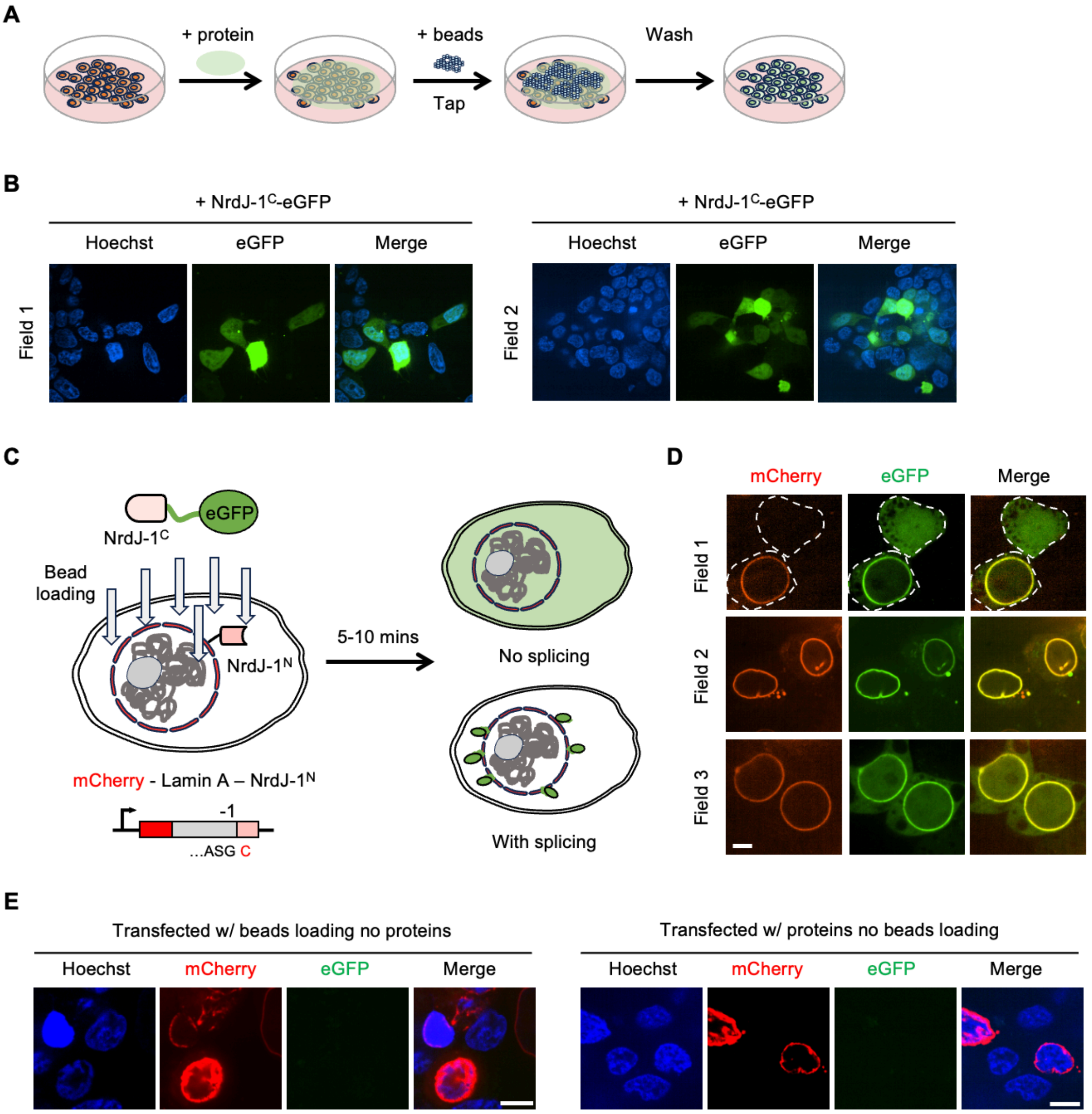


**Supplementary Figure 15. Bead loading delivery of recombinant protein into cultured cells.** (A) Schematic of the bead loading workflow. A solution of purified protein is added onto an adherent monolayer of cultured cells, glass beads sprinkled on top, and the dish is tapped briefly to create transient membrane disruptions. The beads and the excess proteins are then removed by washing multiple times before imaging. (B) Representative fluorescence microscopy images of live HEK293T cells after bead loading with NrdJ-1$^{C}$–eGFP–Flag for 20 mins. Hoechst marks nuclei, eGFP channel shows delivery of NrdJ-1$^{C}$–eGFP–Flag. (C) Schematic of the experimental design: live cell protein trans-splicing (PTS) on nuclear lamina by NrdJ-1. HEK293T cells were transiently transfected with mCherry-Lamin A-NrdJ-1$^{N}$, which localizes to the nuclear-lamina. Recombinant NrdJ-1$^{C}$–eGFP-Flag is delivered to cells via bead loading. In non-transfected cells, no splicing can occur and the eGFP signal therefore remains diffused all over the cell. Conversely, upon PTS the split intein ligates eGFP to Lamin A, producing dual-colored spliced products localized at the nuclear envelope within minutes. The N-extein sequence used in this context for splicing was ASG. (D) Confocal microscopy of live HEK293T cells showing mCherry (red), eGFP (green), and the merge. In the splicingpositive cell, GFP signal forms a ring that co-localized with the mCherry-labeled nuclear lamina (the lower left cell in field 1, and cells in field 2 and 3). For untransfected cells, diffuse GFP signal was observed without PTS occurring (the upper right cell in field 1). Dashed lines outline cell boundaries. Scale bar, 5 μm. (E) Negative controls for protein delivery via bead loading. Representative fluorescence images of HEK293T live cells transiently transfected with mCherry-Lamin A-NrdJ-$1^N$ subjected to bead loading in the absence of recombinant protein (left) or incubated with protein constructs without bead loading (right). Cell nuclei were stained with Hoechst (blue). mCherry (red) and eGFP (green) channels are shown as indicated. No GFP signal was detected under either condition. Scale bar, 10 μm.

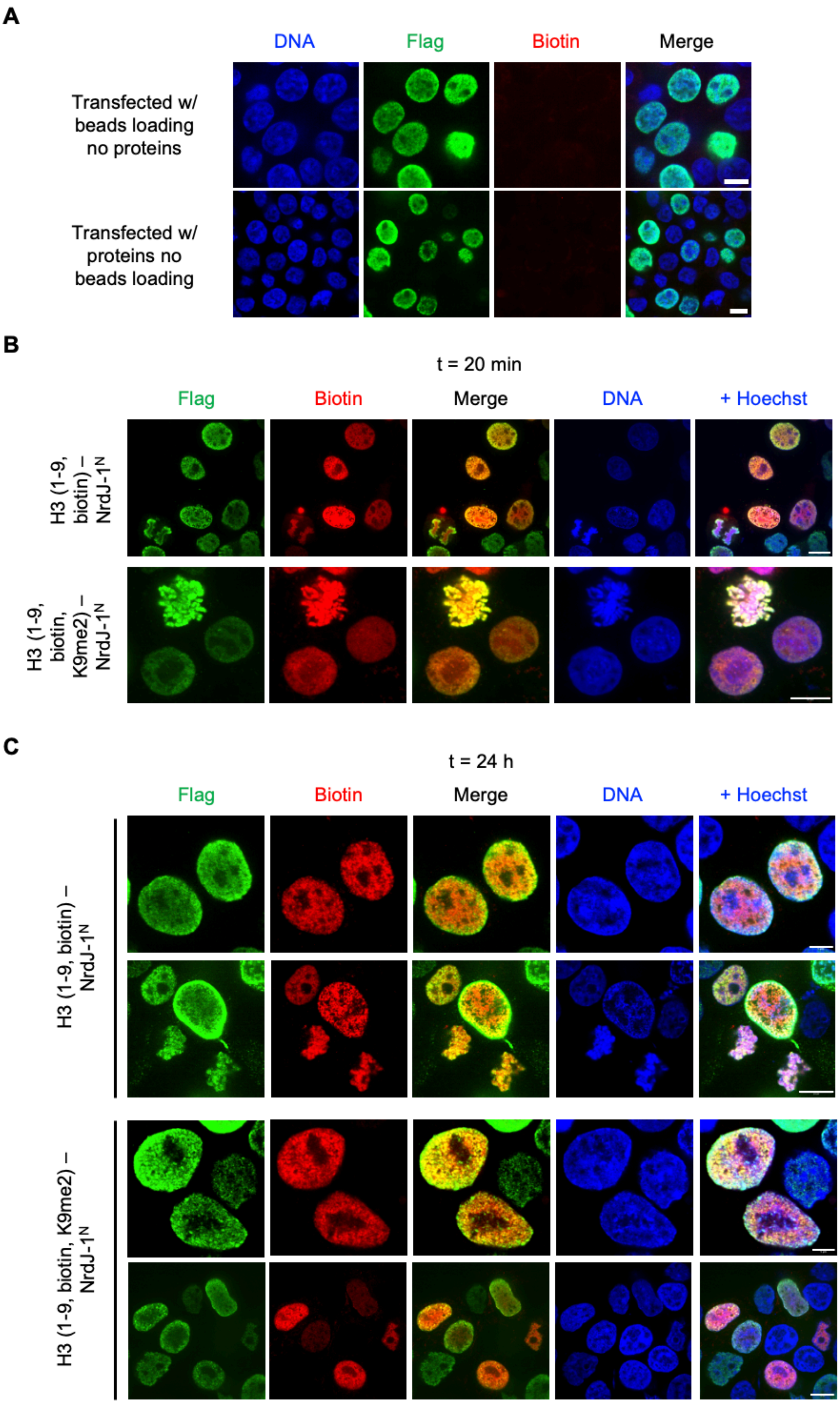
A
DNA
Flag
Biotin
Merge
Transfected w/ beads loading no proteins
Transfected w/ proteins no beads loading
B
t = 20 min
Flag
Biotin
Merge
DNA
+ Hoechst
H3 (1-9, biotin) – NrdJ-1N
H3 (1-9, biotin, K9me2) – NrdJ-1N
C
t = 24 h
Flag
Biotin
Merge
DNA
+ Hoechst
H3 (1-9, biotin) – NrdJ-1N
H3 (1-9, biotin, K9me2) – NrdJ-1N

**Supplementary Figure 16. Confocal microscopy for traceless installation of H3K9 modifications on histone H3 using NrdJ-1** (A) Negative controls for semisynthetic peptide delivery via bead loading. Representative fluorescence images of HEK293T cells transiently transfected with Flag-H3(1–28)-NrdJ-$1^{C}$-H3(10–135) subjected to bead loading in the absence of H3(1–9, biotin)-NrdJ-$1^{N}$ (upper) or incubated with semisynthetic peptide without bead loading (lower). Immunofluorescence staining was performed on fixed cells with anti-Flag (green) and neutravidin (red). DNA is stained with Hoechst (blue). No biotin signal was detected under either condition. Scale bar, 10 μm. (B) Confocal microscopy of HEK293T cells transiently expressing Flag-H3(1–28)-NrdJ-$1^{C}$-H3(10–135). H3(1–9, biotin)-NrdJ-$1^{N}$ and H3(1–9, biotin, K9me2)-NrdJ-$1^{N}$ peptides were introduced into cultured cells by bead loading 24 h post-transfection. Cells were fixed 20 mins post addition of peptides. Immunofluorescence staining was performed as in panel A. Scale bar, 10 μm. (C) Confocal microscopy of HEK293T expressing Flag-H3(1–28)-NrdJ-$1^{C}$-H3(10–135). H3(1–9, biotin)-NrdJ-$1^{N}$ and H3(1–9, biotin, K9me2)-NrdJ-$1^{N}$ peptides were introduced into cultured cells by bead loading 24 h post-transfection, respectively. Cells were fixed 24 hours post addition of peptides. Immunofluorescence staining was performed as described in panel A. Scale bar, upper panel, 5 μm; lower panel, 10 μm.

**Supplementary Table S1.** Kinetics for NrdJ-1 *in vitro* splicing assay.

| | $k_{splice}$ ($s^{-1}$) | $t_{1/2}$ (s) | | $k_{splice}$ ($s^{-1}$) | $t_{1/2}$ (s) |
|---|---|---|---|---|---|
| **WT-1** | 0.02631 | 26.35 | **E2W-1** | 0.006291 | 110.2 |
| **WT-2** | 0.02547 | 27.22 | **E2W-2** | 0.00997 | 69.52 |
| **WT-3** | 0.0284 | 24.41 | **E2W-3** | 0.009565 | 72.46 |
| **C(-1)W-1** | 0.00487 | 142.3 | **E2F-1** | 0.01626 | 42.63 |
| **C(-1)W-2** | 0.007029 | 98.61 | **E2F-2** | 0.02011 | 34.46 |
| **C(-1)W-3** | 0.007739 | 89.56 | **E2F-3** | 0.01498 | 46.27 |
| **C(-1)F-1** | 0.01313 | 52.81 | **E2K-1** | 0.01096 | 63.27 |
| **C(-1)F-2** | 0.01189 | 58.29 | **E2K-2** | 0.01054 | 65.78 |
| **C(-1)F-3** | 0.01384 | 50.07 | **E2K-3** | 0.01516 | 45.74 |
| **C(-1)K-1** | 0.02326 | 29.8 | **E2D-1** | 0.02424 | 28.6 |
| **C(-1)K-2** | 0.0288 | 24.07 | **E2D-2** | 0.02347 | 29.53 |
| **C(-1)E-1** | 0.02393 | 28.97 | **E2D-3** | 0.03226 | 21.49 |
| **C(-1)E-2** | 0.01791 | 38.7 | **E2Q-1** | 0.04024 | 17.23 |
| **C(-1)E-3** | 0.0109 | 63.62 | **E2Q-2** | 0.03487 | 19.88 |
| **C(-1)D-1** | 0.002266 | 305.9 | **E2Q-3** | 0.03514 | 19.73 |
| **C(-1)D-2** | 0.004989 | 138.9 | **E2N-1** | 0.02365 | 29.31 |
| **C(-1)D-3** | 0.001892 | 366.3 | **E2N-2** | 0.02505 | 27.67 |
| **C(-1)Q-1** | 0.06123 | 11.32 | **E2N-3** | 0.0302 | 22.95 |
| **C(-1)Q-2** | 0.0605 | 11.46 | **E2V-1** | 0.0487 | 14.23 |
| **C(-1)Q-3** | 0.05916 | 11.72 | **E2V-2** | 0.01265 | 54.79 |
| **C(-1)N-1** | 0.02967 | 23.36 | **E2V-3** | 0.01366 | 50.76 |
| **C(-1)N-2** | 0.02669 | 25.97 | **E2G-1** | 0.01534 | 45.18 |
| **C(-1)N-3** | 0.03145 | 22.04 | **E2G-2** | 0.02014 | 34.42 |
| **C(-1)V-1** | 0.002569 | 269.8 | **E2G-3** | 0.02336 | 29.67 |
| **C(-1)V-2** | 0.00245 | 282.9 | **E2A-1** | 0.01373 | 50.48 |
| **C(-1)V-3** | 0.00284 | 244.1 | **E2A-2** | 0.01996 | 34.73 |
| **C(-1)G-1** | 0.009614 | 72.09 | **E2A-3** | 0.02513 | 27.58 |
| **C(-1)G-2** | 0.01715 | 40.41 | **E2P-1** | 0.01215 | 57.06 |
| **C(-1)A-1** | 0.009712 | 71.37 | **E2P-2** | 0.02239 | 30.96 |
| **C(-1)A-2** | 0.03247 | 21.34 | **E2P-3** | 0.01848 | 37.52 |
| **C(-1)A-3** | 0.04461 | 15.54 | | | |
| **C(-1)P-1** | 0.00004762 | 14556 | | | |
| **C(-1)P-2** | 2.056E-07 | 3371642 | | | |
| **C(-1)P-3** | 0.002182 | 317.7 | | | |

| | $k_{splice}$ ($s^{-1}$) | $t_{1/2}$ (s) | | $k_{splice}$ ($s^{-1}$) | $t_{1/2}$ (s) |
|---|---|---|---|---|---|
| **P2A-1** | 0.01092 | 63.45 | **Y83A-1** | 0.005727 | 121 |
| **P2A-2** | 0.03037 | 22.82 | **Y83A-2** | 0.005746 | 120.6 |
| **P2A-3** | 0.0285 | 24.32 | **Y83A-3** | 0.008218 | 84.34 |
| **P2E-1** | 0.006942 | 99.85 | **Y89A-1** | 0.001911 | 362.7 |
| **P2E-2** | 0.0106 | 65.37 | **Y89A-2** | 0.004622 | 150 |
| **P2E-3** | 0.008544 | 81.13 | **Y89A-3** | 0.002213 | 313.2 |
| **P2R-1** | 0.01192 | 58.17 | **H39A-1** | 0.002868 | 241.7 |
| **P2R-2** | 0.01854 | 37.39 | **H39A-2** | 0.002554 | 271.4 |
| **P2R-3** | 0.01651 | 41.99 | **H39A-3** | 0.0002716 | 2552 |
| **P2W-1** | 0.01124 | 61.64 | **C76A-1** | 0.02006 | 34.55 |
| **P2W-2** | 0.01847 | 37.54 | **C76A-2** | 0.01465 | 47.31 |
| **P2W-3** | 0.0165 | 42.01 | **C76A-3** | 0.01247 | 55.59 |
| **AAA-1** | 0.01102 | 62.91 | **C76S-1** | 0.01549 | 44.74 |
| **AAA-2** | 0.01271 | 54.52 | **C76S-2** | 0.009415 | 73.62 |
| **AAA-3** | 0.01626 | 42.64 | **C76S-3** | 0.01101 | 62.98 |
| **I3A-1** | 0.02112 | 32.82 | **C76V-1** | 0.006448 | 107.5 |
| **I3A-2** | 0.02223 | 31.18 | **C76V-2** | 0.00969 | 71.53 |
| **I3A-3** | 0.02056 | 33.72 | **C76V-3** | 0.009571 | 72.42 |
| **I3E-1** | 0.01705 | 40.66 | | | |
| **I3E-2** | 0.01622 | 42.73 | | | |
| **I3E-3** | 0.01785 | 38.84 | | | |
| **I3R-1** | 0.01443 | 48.04 | | | |
| **I3R-2** | 0.01933 | 35.86 | | | |
| **I3R-3** | 0.01892 | 36.63 | | | |
| **I3W-1** | 0.009057 | 76.53 | | | |
| **I3W-2** | 0.01587 | 43.69 | | | |
| **I3W-3** | 0.02874 | 24.12 | | | |
| **SAA-1** | 0.01603 | 43.24 | | | |
| **SAA-2** | 0.009911 | 69.94 | | | |
| **SAA-3** | 0.01539 | 45.03 | | | |

### Materials and Methods

*Antibody*

| Antibody (Host) | Usage and Dilution | Source | Catalog Number |
|---|---|---|---|
| anti-FLAG (mouse) | Western blot – 1:5000<br>Immunofluorescence – 1:2000 | Sigma | F1804 |
| anti-H3 (rabbit) | Western blot – 1:5000 | Abcam | Ab1791 |
| anti- H3Q5ser (rabbit) | Western blot – 1:2000 | Gift from Ian Maze lab | N/A |
| anti-H3K9ac (rabbit) | Western blot – 1:2000 | Abcam | Ab4441 |
| anti-H3K9me2 (mouse) | Western blot – 1:2000 | Abcam | Ab1220 |
| IRDye® 680RD –<br>anti-Mouse IgG (goat) | Western blot – 1:10000 | LI-COR | 926-68070 |
| IRDye® 800CW –<br>anti-Rabbit IgG (goat) | Western blot – 1:10000 | LI-COR | 926-32211 |
| IRDye® 800CW Streptavidin | Western blot – 1:10000 | LI-COR | 926-32230 |
| anti-Mouse IgG (goat) –<br>Alexa Fluor™ 488 | Immunofluorescence – 1:2000 | ThermoFisher | A-11001 |
| NeutrAvidin™, Rhodamine Red™-X conjugate | Immunofluorescence – 1:1000 | ThermoFisher | A6378 |

*DNA constructs and molecular cloning*

| Name | Fusion components | Note |
|---|---|---|
| MBP-NrdJ-$1^N$ in pET | His-SUMO-HA-MBP-NrdJ-$1^N$ | NGTNPC as N-extein, used in ITC, extein dependence and Cys mutation, and accelerator experiments |
| NrdJ-$1^C$-SUMO (N145A) in pET | His-SUMO- NrdJ-$1^C$(N145A)-SUMO | For ITC experiment |
| NrdJ-$1^C$-eGFP in pET | His-SUMO-NrdJ-$1^C$-eGFP-Flag | SEIVL as C-extein, used in extein dependence, Cye mutation, and accelerator experiments |
| NrdJ-$1^N$ (C76A) in pET | His-SUMO-NrdJ-$1^N$ (C76A) | Semi-synthesis of H3(1-9)-NrdJ-$1^N$ (C76A) |
| NrdJ-$1^C$-H3.1(10-130) in pET | NrdJ-$1^C$-H3.1(10-130) | Octamer assembly for *in vitro* splicing |
| Flag-H3(1-28)-NrdJ-$1^C$-H3.1 in pCMV | Flag-H3(1-28)-NrdJ-$1^C$-H3.1 | Splicing in isolated nuclei and in live cells for histone H3 |
| mCherry-LMNA-NrdJ-$1^N$ in pCMV | Flag-mCherry-progerin (NrdJ-$1^N$) | NrdJ-$1^N$ inserted into the C-terminal of the progerin protein, for nuclear envelope splicing experiments |

All constructs used in this study were generated using standard molecular cloning procedures (Gibson assembly, restriction cloning, site-directed mutagenesis) and confirmed by Sanger sequencing (Genewiz) or full plasmid seuqencing (Plasmidsaurus). Plasmid maps and protein sequences for all constructs used in this study are given below (pages S35-S42).

Oligonucleotide primers for cloning were purchased from Integrated DNA Technologies (IDT) or MilliporeSigma. gBlock® Gene Fragments were synthesized by IDT. PrimeSTAR® HS DNA Polymerase (premix, Takara R040A) was used for PCR. Gibson Assembly Master Mix was purchased from New England Biolabs (E2611L). BL21(DE3) chemically competent E. coli cells and Subcloning Efficiency DH5α competent *E. coli* cells were generated in-house from cells purchased from Invitrogen. DNA purification kits for plasmid purification were purchased from QIAGEN. PCR Purification & Gel extract columns were purchased from Thomas Scientific.

*General Equipment*

Analytical RP-HPLC was performed on an Agilent 1260 Infinity system with a Vydac C18 column (5 µm, 4.6 x 150 mm) at a flow rate of 1 mL/min. Semi-preparative RP-HPLC was performed on an Agilent 1260 Infinity system with a Zorbax 300SB-C18 column (5 µm, 9.4 x 250 mm) at a flow rate of 4 mL/min. Large scale peptide purification was performed with preparative Waters RP-HPLC system with a 2535 Quartenary Gradient Module employing an XBridge Peptide BEH C18 OBD Prep Column, 300Å (10 µm, 19 × 250 mm) (Waters) at a flow rate of 20 mL/min. All runs used 0.1 % TFA (trifluoroacetic acid) in water (solvent A) and 90 % acetonitrile in water with 0.1 % TFA (solvent B) as the mobile phases. Unless otherwise stated, peptides and proteins were analyzed using the following gradients: 0% B for 2 minutes followed by 0-100% B over 30 or 40 minutes. Electrospray ionization mass spectrometric analysis (ESI-MS) was performed on a Bruker Daltonics MicroTOF-Q II mass spectrometer. Lyophilization of samples after HPLC was performed on a Millrock Technology MD85 lyophilizer (Kingston, NY). Cell disruption (sonication) was performed with an Ultrasonic Liquid Processor Sonic Dismembrator (ThermoFisher). All size-exclusion chromatography (SEC) was performed on a GE Healthcare (Chicago, IL) AKTA PV-908 FPLC system or a Cytiva (Marlborough, MA) AKTA Go protein purification system with a Superdex S200 increase 10/300 column. SDS-PAGE gels and western blots were imaged on a LI-COR Odyssey Photoimager (Lincoln, NE) or a Cytiva Amersham ImageQuant 800 instrument (Malborough, MA).

*Protein expression and purification*

For all the recombinant proteins except histones, chemically competent *E. coli* BL21(DE3) cells were heat-shock transformed with a pET vector carrying the gene cassette for the protein of interest with related extein sequences. Cells were grown overnight at 37 °C in 6 mL LB medium supplemented with 50 µg/mL kanamycin. This overnight culture was used to inoculate an expression culture of 1 L LB medium supplemented with 50 µg/mL kanamycin. In general, the expression culture was incubated at 37 °C until it reached an OD600 of 0.4-0.6, whereafter it was cooled for 20 min at 18 °C. Protein expression was then induced by the addition of 0.1 mL 1 M IPTG and the culture left overnight at 18 °C. The overnight cultures were harvested by centrifugation at 3,500 g, 4 °C for 30 min and suspended in lysis buffer containing 20 mL 50 mM $NaH_2PO_4$ (pH 8.0), 300 mM NaCl, 20 mM imidazole, supplemented with 1 mM dithiothreitol (DTT) and 1 mM phenylmethylsulfonyl fluoride (PMSF). Cell suspensions were then either stored at –80 °C until further use or used directly. Soluble protein was extracted by subjecting the cell suspension to sonication using a duty cycle of 20s on, 30s off, at 30% amplitude for 4 min 40 s while cooled on an ice bath, after which a cleared lysate was produced by centrifugation at 35,000 g, 4 °C for 20 min. The cleared lysate was passed through a pre-equilibrated Nickel-nitrilotriacetic acid (Ni-NTA) resin (Thermo Scientific, 4 mL resin slurry per liter culture) and the flow-through discarded. The column was then washed with 50 mL lysis buffer, before the protein was eluted using 6 mL lysis buffer supplemented with 250 mM imidazole. $His_6$-SUMO tagged proteins were treated overnight with $His_6$-Ulp1 protease while being dialyzed against 4 L lysis buffer supplemented with 1 mM DTT. The dialyzed sample was then passed through a preequilibrated Ni-NTA column to remove any cleaved $His_6$-SUMO tag and $His_6$-Ulp1 protease. The flow-through and an additional 6 mL lysis buffer passed through the column was collected, combined, and concentrated to 0.5 mL.

For proteins used in *in vitro* splicing reactions, the concentrated sample was filtered through a 0.22 µm spin filter. Size exclusion chromatography was performed at 4 °C at a flowrate of 0.3 mL/min on an ÄKTA Fast Performance Liquid Chromatography (GE Healthcare) system using a Superdex 200 10/300 GL (Cytiva Life Sciences) column with 100 mM $NaH_2PO_4$ (pH 7.2), 150 mM NaCl, 1 mM ethylenediaminetetraacetic acid (EDTA), 10% glycerol and 1 mM DTT as the eluent. For proteins used isothermal titration calorimetry, buffer without glycerol and DTT were used. Pure fractions were identified by SDS-PAGE (10% Bis-Tris acrylamide gel), validated by analytical RP-HPLC, and the protein mass confirmed by ESI-TOF MS. Pure fractions were aliquoted and flash-frozen in liquid nitrogen before being stored at -80 °C until further use. For protein used in semisynthesis, the concentrated sample was filtered with a 0.45 um syringe filter and purified by preparative C-18 RP-HPLC, using a gradient 20-60% B gradient over 60 min. Purified proteins were lyophilized and analyzed by analytical RP-HPLC and ESI-MS. Analytical data and protein sequences for all proteins is shown in Supplementary Figures.

*Purification of histones*

The purification of histones was adapted from a previously reported protocol[42]. Unmodified recombinant human histones (H2A, Uniprot ID: Q6FI13; H2B, Uniprot ID: O60814; H4, Uniprot ID: P62805), and NrdJ-1$^{C}$-H3.1 (10-135) were produced in and purified from *E. coli*. In brief, BL21 Rosetta (DE3) cells were transfected with histone expression plasmids and grown in LB medium at 37 °C until they reached an OD600 of 0.6. Protein expression was induced by the addition of 0.5 mM IPTG for 2–4 h at 37 °C. Cells were harvested by centrifugation at 4,000 g for 30 min at 4 °C. Cell pellets were resuspended in 10 ml of cold lysis buffer (50 mM Tris, 100 mM NaCl, 1 mM EDTA, 5 mM 2-mercaptoethanol, 1mM PMSF, pH 7.6 at 4 °C). Cells were lysed via sonication (20% Amplitude, 30 seconds ON, 30 seconds OFF for a total on time of 3 mins 30 seconds) and the resulting suspension was centrifuged at 35,000 g for 30 min at 4 °C. The inclusion body pellet was washed twice with cold lysis buffer containing 1% Triton-X 100 and once without detergent. Inclusion body pellets were resuspended in 20 ml inclusion body resuspension buffer (6 M guanidine hydrochloride, 20 mM Tris, 1 mM EDTA, 100 mM NaCl, 5 mM 2-mercaptoethanol, pH 7.5 at 4 °C) per liter of culture, and nutated at 4 °C for 2 h. Resuspensions were then centrifuged at 35,000 g for 30 min at 4 °C. The supernatants were transferred to the ultracentrifuge for 30 minutes at 50,000 RPM at 4 °C after acidifying using HCl to pH towards 2 or below. The histone-containing supernatant solution was then filtered with a 0.45 um syringe filter and the histones were purified by preparative C-18 RP-HPLC using a gradient 20-60% B gradient over 60 min (for H2A, H2B, and H4) or a 30-70% B gradient over 60 minutes (for H3). Purified histones were lyophilized and analyzed by analytical RP-HPLC and ESI-MS.

*Histone octamer formation*

Histone octamers were formed using a previously reported protocol[42]. Lyophilized purified histones were dissolved in histone unfolding buffer (6 M guanidine hydrochloride, 20 mM Tris, 5 mM DTT, pH 7.5 at 4 °C) and combined in the following ratios (0.0275 nmol each of histones H2A, H2B, and 0.02275 nmol each of histones NrdJ-1C-H3.1 (10-135), and H4). Use of excess of H2A and H2B minimizes the presence of H3-H4 tetramers (which do not fully resolve from the desired octamers on the FPLC) in the preparation. The total histone concentration was adjusted to 1 mg/ml, and the mixtures were placed in Slide-A-Lyzer MINI dialysis devices (3.5 kDa MW cutoff, ThermoFisher Scientific) and dialysed at 4 °C against 3 × 400 ml of octamer refolding buffer (2 M NaCl, 10 mM Tris, 0.5 mM EDTA, 1 mM DTT, pH 7.8 at 4 °C) for at least 4 h for each step, with one dialysis step overnight. The mixtures were then filtered and further purified by size exclusion chromatography using an S200 increase 10/300 gel filtration column with octamer refolding buffer as the mobile phase. Fractions were analyzed by SDS-PAGE. The octamer fractions were transferred to fresh microcentrifuge tubes, and 50% (v/v) glycerol was added. Octamer concentrations were measured by UV spectroscopy at 280 nm, and stored at 4 °C.

*Isothermal titration calorimetry (ITC)*

ITC was performed using a MicroCal PEAQ-ITC instrument (GE Healthcare). MBP–NrdJ-$1^N$ and NrdJ-$1^C$–SUMO constructs carrying inactivating mutations (C1A in the N-intein and N145A in the C-intein) were expressed and purified as described in the previous section. Both proteins were concentrated and dialyzed overnight into ITC buffer (100 mM $NaH_2PO_4$, 150 mM NaCl, pH 7.35) using dialysis buttons (Slide-A-Lyzer Mini Dialysis unit, 3.5 MWCO, ThermoScientific). For ITC experiments, NrdJ-$1^C$–SUMO (110 µM) was loaded into the syringe and titrated into the sample cell containing MBP–NrdJ-$1^N$ (7 µM) in a total volume of ~300 µl. Approximately 70 µl of NrdJ-$1^C$–SUMO solution was automatically injected in a series of aliquots at 25°C. Data were processed and analyzed using the MicroCal PEAQ-ITC analysis software.

*In vitro splicing assays*

The *in vitro* splicing procedure and analysis was adapted from a previously reported protocol[15,16,43]. For general intein splicing, N- and C-inteins (1 µM $Int^N$, 1µM $Int^C$) were individually preincubated in splicing buffer (100 mM sodium phosphates, 150 mM NaCl, 1 mM EDTA, pH 7.2, 10% glycerol) with 1 mM DTT for 15 minutes at 37 °C. For in vitro splicing on assembled octamer, 2 µM of NrdJ-$1^C$ octamer and 5 µM of various H3 (1-9)-NrdJ-$1^N$ constructs were used. Splicing reactions were carried out at 37 °C. Splicing was initiated by mixing equal volumes of N- and C- inteins with aliquots removed at the indicated times and quenched by the addition of 4X loading dye (160 mM Tris, 40% glycerol, 4% SDS, 0.08% Bromophenol Blue, 8 % BME). Samples were analyzed by SDS-PAGE (16% Tris-glycine gels for PTS involving histones and 10 % Bis-Tris acrylamide gels for all the other PTS experiments) and starting materials and product bands quantified by densitometry using ImageJ.

To determine the splicing rates of trans-splicing reactions, the data was fit to the first order rate equation using the GraphPad Prism software.

$[P](t) = [P]max (1-e^{-kt})$

Where [P] is the normalized intensity of product, [P]max is this value at t=∞ (the reaction plateau), and k is the rate constant ($s^{-1}$). The mean and standard deviation are calculated by GraphPad Prism. Kinetics of for all replicates are reported in supplementary table 1.

*Peptide synthesis*

Histone H3 peptides were synthesized by manual Fmoc-SPPS on ChemMatrix Trityl-OH resin, preloaded with a hydrazine linker. The solid phase synthesis cycle for non-modified amino acids included: i) Fmoc deprotection with 20% piperidine in DMF (15 min) and ii) coupling of 5 equivalents of each amino acid in DMF with 5 equivalents of (7-Azabenzotriazol-1-yloxy)tripyrrolidinophosphonium hexafluorophosphate (PyAOP) and 10 equivalents diisopropylethylamine (DIPEA) for 40 min. For modified amino acids, 2 equivalents of each amino acid (Fmoc-Lys(ac)-OH, Fmoc-Lys(biotin)-OH, Fmoc-Lys(me2)-OH, Fmoc-Glu(OAll)-OH) in DMF was used during coupling instead. The coupling step was repeated twice for all amino acids. To generate H3 peptides carrying H3Q5ser, Fmoc-Glu(OAll)-OH was incorporated at residue 5 to allow subsequent serotonylation after allyl deprotection[34]. Fully assembled resin-bound peptides were subjected to allyl deprotection by treatment with 0.2 equivalents of $Pd(PPh_3)_4$ and 10 equivalents of DIPEA in DCM, applied twice for 20 min each. The resin was then extensively washed with DCM and DMF to remove residual palladium. Serotonylation was performed by treating the resin with 1.1 equivalents of PyAOP, 5 equivalents of DIPEA, and 2 equivalents of serotonin hydrochloride in DMF, applied as double couplings of 30 min each. Finally, peptide cleavage and side-chain deprotection were carried out with a cleavage cocktail (95% trifluoracetic acid (TFA), 2.5% triisopropylsilane (TIPS), 2.5% $H_2O$) for 2 hours. Cold diethyl ether was added, and the peptides were precipitated and collected by centrifugation at 4,000g for 10 minutes. The peptides were characterized by analytical C18 RP-HPLC and ESI-MS.

*Generation of semisythetic products*

4-Mercaptophenylacetic acid (MPAA) thioester generation: crude peptides obtained from SPPS were converted into the corresponding MPAA thioesters following a previously reported procedure[44]. Briefly, peptide hydrazides were dissolved in conversion buffer (6 M guanidine HCl, 100 mM sodium phosphate, pH 3.0) with 100 nM MPAA and 2.5 mM acetyl acetone (acac) to achieve final concentrations of 1 mM peptide (calculated from crude mass). The pH of the reaction mixture was adjusted to 3.0 and shaken vigorously overnight at room temperature. Reaction mixtures were then diluted with HPLC solvent A, acidified to pH 2.0, and purified by preparative RP-HPLC (0% B for 5 min, followed by a 20–60% B gradient over

40 min). The purified products were identified by ESI-MS, collected, and lyophilized for storage.

Native chemical ligation: the lyophilized MPAA-thioester fragment (1 mM) and NrdJ-$1^N$ (C76A, 0.5 mM) were mixed at room temperature in ligation buffer (6M guanidine HCl, 100 mM sodium phosphate buffer, pH 7.0, 50 mM Tris(2-carboxyethyl)phosphine hydrochloride (TCEP), 100 mM MPAA) and allowed to react overnight. The reaction mixtures were then diluted with HPLC Solvent A, acidified to pH 2 and purified by semi-preparative RP-HPLC (0% B for 5 minutes, then 20-60% B over 40 minutes). The products were identified by ESI-MS, collected and lyophilized. The lyophilized products were pooled and dissolved in denaturing buffer (6M Urea, 100 mM sodium phosphate buffer, pH 7.4, 1 mM DTT) and then dialyzed in a stepwise fashion (4M Urea, 2M Urea, 0M Urea with 100 mM sodium phosphate buffer, pH 7.4, 1 mM DTT) into PBS buffer (pH 7.4). Glycerol was added to a final concentration of 10%, after which the products were aliquoted, flash-frozen in liquid nitrogen and stored at –80 °C until use.

*Cell culture and transfection*

HEK293T (purchased from ATCC) were cultured in Dulbecco's Modified Eagle Medium (DMEM, Thermo Fisher 11995-005), supplemented with 10% v/v FBS (Atlanta Biologicals S12450H), 100 U/mL penicillin, 100 µg/mL streptomycin (ThermoFisher 15140-122), and 2 mM L-Glutamine (Thermo Fisher 25030-081). Cells were maintained in an incubator at 37 °C with 5% $CO_2$. For beads loading and imaging experiments, tissue culture plates were pre-coated with 0.01 % poly-L-lysine solution (Sigma P8920).

Transfection for transient protein expression in cells was performed using Lipofectamine™ 2000 Transfection Reagent (Thermo Fisher 11668-019) following the manufacturer’s instructions. For *in nucleo* splicing assays, the cells were expanded into 10 cm plates and transfected with a plasmid encoding FLAG-$H3_{1\text{-}28}$-NrdJ-$1^C$-H3.1(10-135). After 24 h, the plates were harvested. The cell pellets were flash frozen and stored at –80 °C for further processing. For live cell protein trans splicing (PTS), the cells were cultured in a 35 mm glass-bottom petri dish (MatTek P35G-1.5-14-C) plates pre-coated with 0.01 % poly-L-lysine solution. At about 50% confluency, the cells are transfected with the plasmid encoding FLAG-$H3_{1\text{-}28}$-NrdJ-$1^C$-H3.1(10-135) or FLAG-mCherry-LMNA-NrdJ-$1^N$. After 24 h, the cells were ready for further PTS experiments.

*PTS in nucleo*

The *in nucleo* splicing procedure was adapted from a previously reported protocol[35]. The frozen transfected pellet was lysed under hypotonic conditions using 1 mL of RSB buffer (10 mM Tris pH 7.4, 15 mM NaCl, 1.5 mM $MgCl_2$, Roche cOmplete EDTA-free protease inhibitors) for 10 min on ice. The crude nuclei were isolated by centrifugation (Note: all centrifugations were conducted at 500 g for 5 min at 4 °C unless noted otherwise), resuspended in 1 ml RSB buffer, and homogenized for using ten strokes of a loose pestle Dounce homogenizer. The isolated nuclei were then pelleted by centrifugation and resuspended in 500 µL of PTS buffer (20 mM HEPES, 1.5 mM $MgCl_2$, 150 mM KCl, Roche cOmplete EDTA-free protease inhibitors, pH 7.4). The nuclei were centrifuged and washed again with 500 µl of PTS buffer. Finally, the pelleted nuclei were resuspended in 300 µl of PTS buffer per 10 cm dish of transfected HEK293T cells. An indicated amount of modified $H3_{1-9}$-NrdJ-$1^N$(C76A) was added to each nuclei aliquot. The reactions were incubated at 37 °C for 1 hour. Next, the treated nuclei were dissolved in 1x SDS sample loading buffer (100 mM Tris-Cl, pH 6.8, 3% SDS, 15% Glycerol, 2.25% β-mercaptoethanol, 0.015% bromophenol blue). Samples were boiled at 98 °C for 30 mins and were subsequently analyzed by western blot analysis.

*Western blot*

Samples were resolved on a bis-tris or tris-glycine polyacrylamide gel. For Western blotting, the gel was used for a transfer reaction onto a nitrocellulose membrane, which was subsequently blocked with TBS-T (25 mM Tris, 150 mM NaCl, 0.1% v/v Tween-20, pH 7.7) supplemented with 4% w/v skimmed milk powder for 30 mins at RT. The membrane was washed 3 x 5 min with TBS-T and incubated with primary antibodies (diluted in TBS-T containing 4% w/v Bovine Serum Albumin (BSA)) at a specified dilution on an orbital shaker for 2 hr at room temperature or alternatively overnight at 4 °C. After washing 3 x 5 min with TBS-T, the IRDye secondary antibody (1:10000 dilution in TBS-T) or alternatively IRDye Streptavidin (1:10000 dilution in TBS-T) were applied for 30 min at room temperature, before imaging on a Li-Cor Odyssey imager (Li-Cor).

*Bead loading*

The bead loading procedure was adapted from a previously reported protocol[40]. In brief, 5 ml of glass beads (⩽106 µm; Sigma-Aldrich, G4649) were washed in a 50 ml conical tube with 25 ml NaOH for 2 h, followed by five washes with 25 ml Milli-Q water until neutral pH was confirmed using pH test strips. The beads were then washed three times with 100% ethanol, air-dried overnight in a 10 cm Petri dish (with the lid open) inside a sterile cell-culture hood, and finally UV-sterilized for 15 min in the cell-culture hood.

HEK293T cells were transfected and plated in 35 mm glass-bottom Petri dishes as described earlier and grown to 80–90% confluency 24 hours after transfection. Prior to bead loading, cell medium was removed, and the cells were washed 1–2 times with sterile DPBS. A total of 6–8 µl of 20 µM protein solution in splicing buffer (100 mM $NaH_2PO_4$, pH 7.2; 150 mM NaCl; 1 mM EDTA; 1 mM DTT; 10% v/v glycerol) was added directly to the glass coverslip portion of the dish and allowed to spread for 5 s. A monolayer of glass beads was then gently dispersed on top of the protein solution using a 100 µm diameter sterile cell strainer, applied by lightly tapping 8-10 times to ensure complete coverage of the cells in the glass-bottom well. After bead application, culture medium was carefully added back to the dish by pipetting slowly along the plastic edge of the chamber to avoid disturbing the cells. Cells were then washed 1–

2 times using either DPBS or culture media to remove excess beads and protein solution, and floating beads were aspirated without disturbing the adherent cells. Pre-warmed complete medium was then added, and the cells were incubated for the indicated PTS reaction times. Before imaging, cells were washed once with culture medium to remove any residual beads.

*Immunofluorescence*

Cells were cultured in either a 35 mm glass-bottom petri dish (MatTek P35G-1.5-14-C) or a 24-well glass-bottom plate (Cellvis P24-1.5H-N). Cells were transfected and treated as described above. Subsequently, cells were fixed with 4% formaldehyde for 10 min, then permeabilized with 0.5% Triton in PBS for 15 min at room temperature. 3% BSA in PBS was used for blocking for at least 1 hour at room temperature or overnight at 4 °C. For experiments that only need biotin staining, cells were incubated with NeutrAvidin™-Rhodamine Red™-X (2 µg/mL, ThermoFisher A6378) and Hoechst 33342 (10 µg/mL) for 1 h at RT, washed 3 times with PBS, and imaged using a confocal microscope. When staining with a primary antibody was also needed, the antibody was incubated with the cells overnight at 4 °C, followed by staining with a secondary antibody, NeutrAvidin™-Rhodamine Red™-X, and Hoechst 33342 for 1 h at RT. Cells were washed 3 times with PBS and imaged using a confocal microscope.

*Confocal microscopy*

A 488 nm laser was used for the excitation of eGFP, 405 nm for the excitation of Hoechst 33342, and 561 nm for Rhodamine Red/mCherry excitation. Imaging was performed using a Nikon CSU-21 spinning disk confocal microscope equipped with NIS Elements software (version 5.2), a 60× Plan Apo λ oil-immersion objective, and a Hamamatsu ORCA-Flash4.0 sCMOS camera.

**DNA constructs and protein sequences**

MBP-NrdJ-1$^{N}$ in pEET

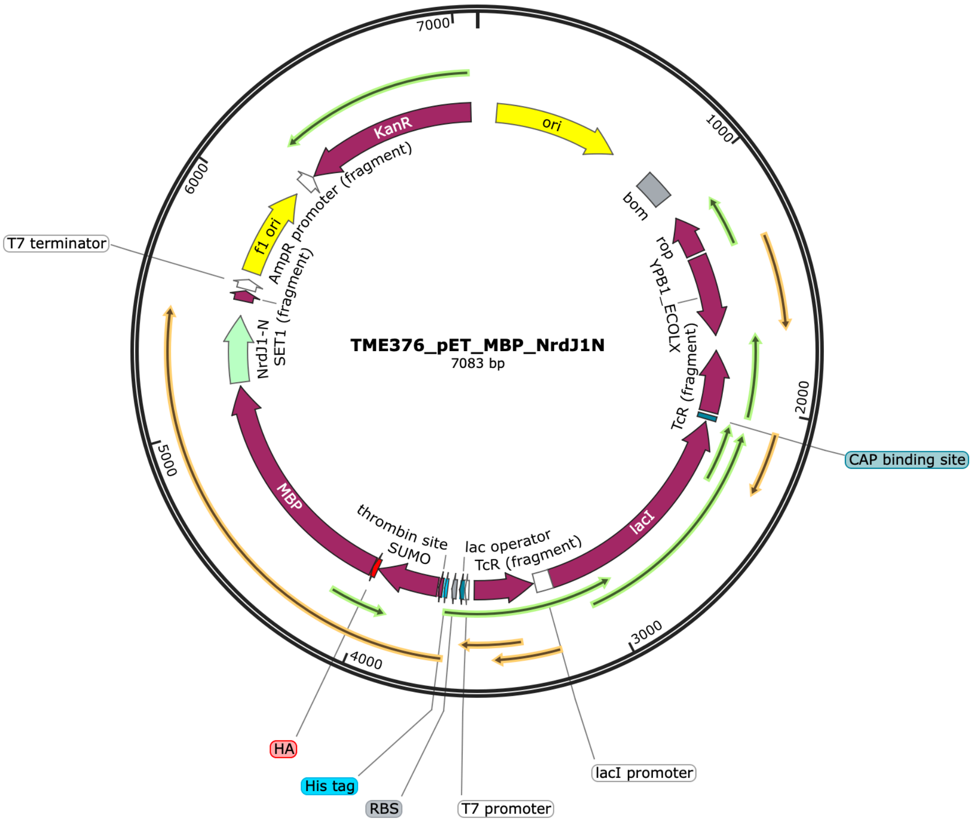


Expressed protein sequence (His-SUMO-HA-MBP-NrdJ-1$^{N}$):

MGSSHHHHHHGSGLVPRGSASMSDSEVNQEAKPEVKPEVKPETHINLKVSDGSSEIFF
KIKKTTPLRRLMEAFAKRQGKEMDSLRFLYDGIRIQADQTPEDLDMEDNDIIEAHREQ
IGGYPYDVPDYAKIEEGKLVIWINGDKGYNGLAEVGKKFEKDTGIKVTVEHPDKLEE
KFPQVAATGDGPDIIFWAHDRFGGYAQSGLLAEITPDKAFQDKLYPFTWDAVRYNGK
LIAYPIAVEALSLIYNKDLLPNPPKTWEEIPALDKELKAKGKSALMFNLQEPYFTWPLI
AADGGYAFKYENGKYDIKDVGVDNAGAKAGLTFLVDLIKNKHMNADTDYSIAEAA
FNKGETAMTINGPWAWSNIDTSKVNYGVTVLPTFKGQPSKPFVGVLSAGINAASPNK
ELAKEFLENYLLTDEGLEAVNKDKPLGAVALKSYEEELAKDPRIAATMENAQKGEIM
PNIPQMSAFWYAVRTAVINAASGRQTVDEAPKDAQTNGTNPCCLVGSSEIITRNYGKT
TIKEVVEIFDNDKNIQVLAFNTHTDNIEWAPIKAAQLTRPNAELVELEIDTLHGVKTIR
CTPDHPVYTKNRGYVRADELTDDDELVVAI*

(Underlined: mutated residues during the studies)

NrdJ-1$^{C}$-SUMO (N145A) in pET

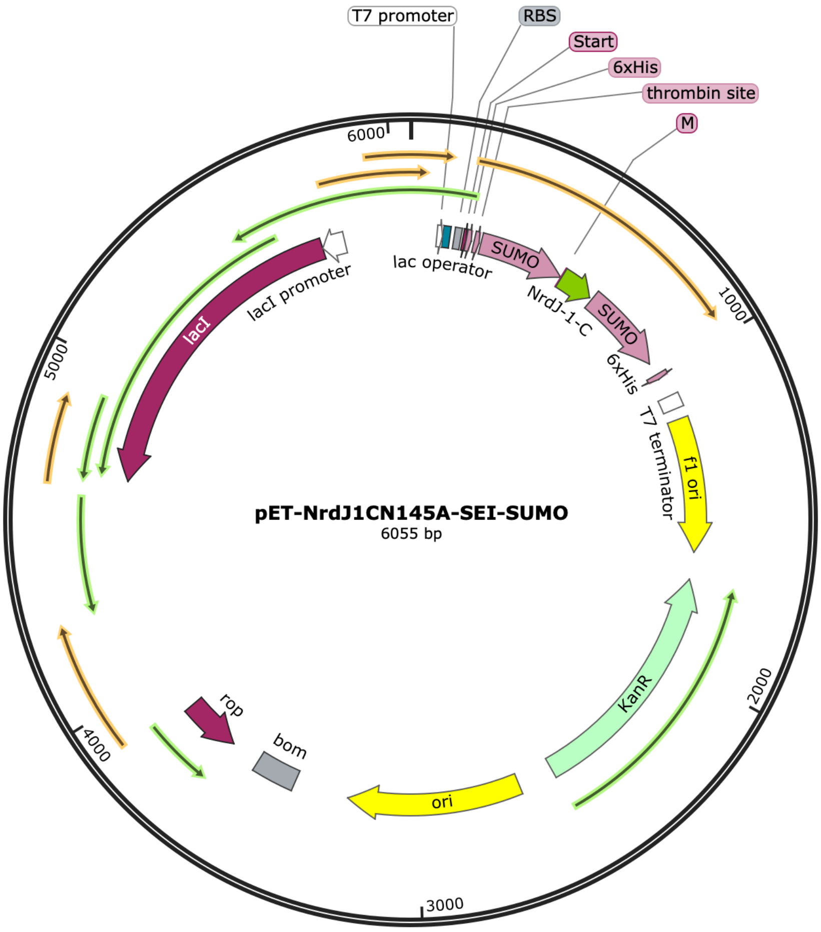


Expressed protein sequence (His-SUMO- NrdJ-1$^{C}$(N145A)-SUMO):

MGSSHHHHHHGSGLVPRGSASMSDSEVNQEAKPEVKPEVKPETHINLKVSDGSSEIF
FKIKKTTPLRRLMEAFAKRQGKEMDSLRFLYDGIRIQADQTPEDLDMEDNDIIEAHRE
QIGGMEAKTYIGKLKSRKIVSNEDTYDIQTSTHNFFANDILVHASEIMSDSEVNQEAK
PEVKPEVKPETHINLKVSDGSSEIFFKIKKTTPLRRLMEAFAKRQGKEMDSLRFLYDGI
RIQADQTPEDLDMEDNDIIEAHREQIGG*

NrdJ-1C-eGFP in pET

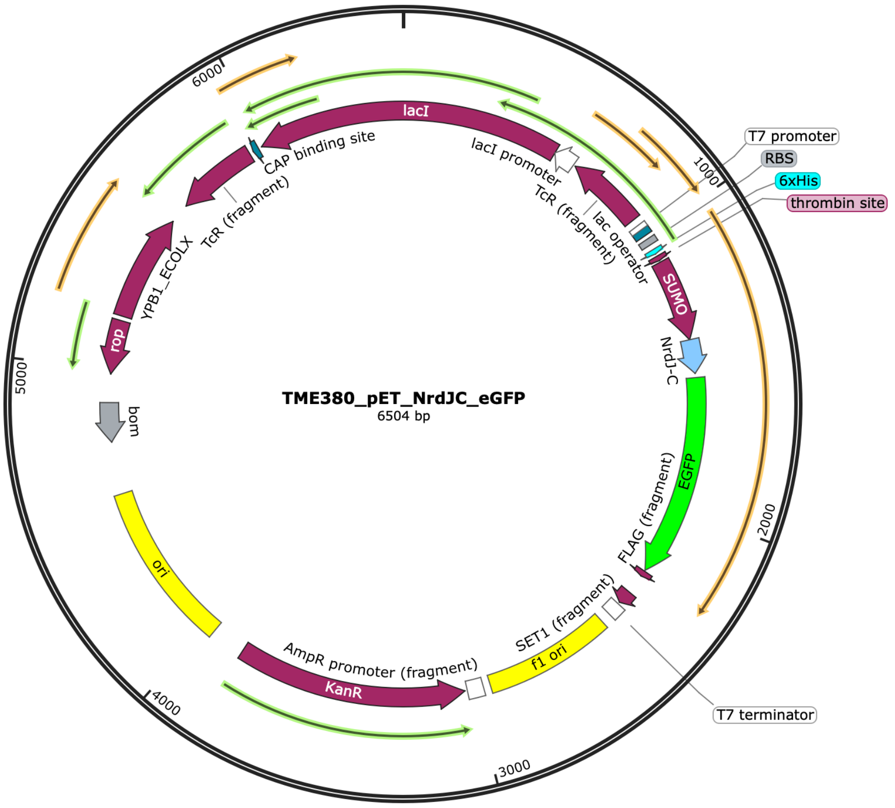


Expressed protein sequence (His-SUMO-NrdJ-1C-eGFP-Flag):

MGSSHHHHHHGSGLVPRGSASMSDSEVNQEAKPEVKPEVKPETHINLKVSDGSSEIF
FKIKKTTPLRRLMEAFAKRQGKEMDSLRFLYDGIRIQADQTPEDLDMEDNDIIEAHRE
QIGGMEAKTYIGKLKSRKIVSNEDTYDIQTSTHNFFANDILVHNSEIVLMVSKGEELFT
GVVPILVELDGDVNGHKFSVSGEGEGDATYGKLTLKFICTTGKLPVPWPTLVTTLTYG
VQCFSRYPDHMKQHDFFKSAMPEGYVQERTIFFKDDGNYKTRAEVKFEGDTLVNRIE
LKGIDFKEDGNILGHKLEYNYNSHNVYIMADKQKNGIKVNFKIRHNIEDGSVQLADH
YQQNTPIGDGPVLLPDNHYLSTQSALSKDPNEKRDHMVLLEFVTAAGITLGMDELYK
DYKDDDDK*

(Underlined: mutated residues during the studies)

NrdJ-1$^{N}$ (C76A) in pET

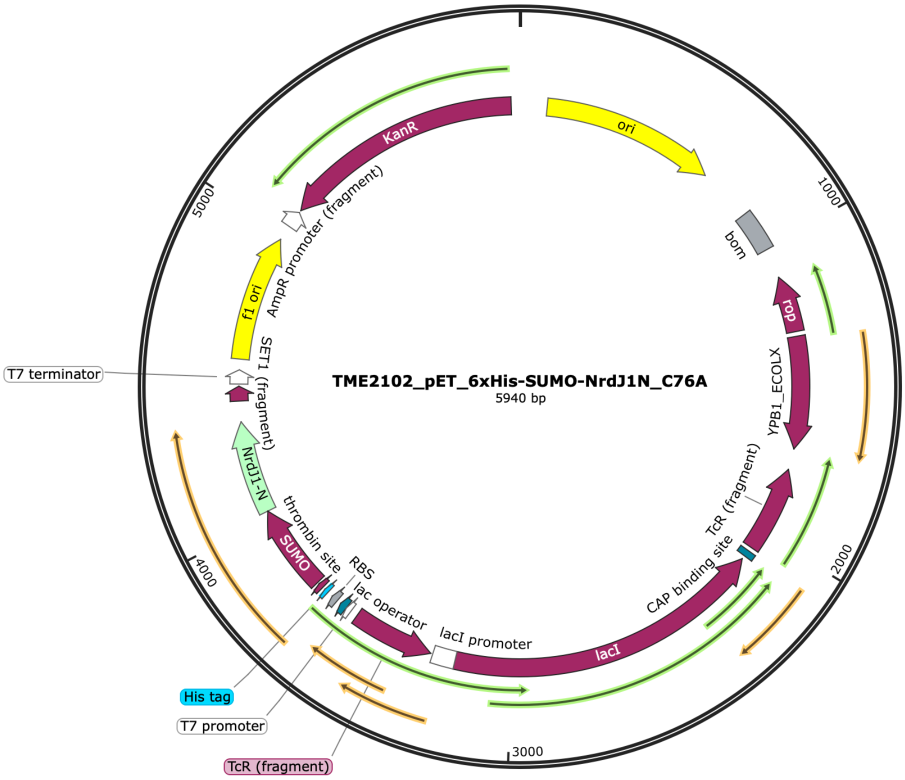


Expressed protein sequence (His-SUMO-NrdJ-1$^{N}$ (C76A)):

MGSSHHHHHGSGLVPRGSASMSDSEVNQEAKPEVKPEVKPETHINLKVSDGSSEIFF
KIKKTTPLRRLMEAFAKRQGKEMDSLRFLYDGIRIQADQTPEDLDMEDNDIIEAHREQ
IGGCLVGSSEIITRNYGKTTIKEVVEIFDNDKNIQVLAFNTHTDNIEWAPIKAAQLTRPN
AELVELEIDTLHGVKTIRATPDHPVYTKNRGYVRADELTDDDELVVAI*

NrdJ-1^C-H3.1(10-130) in pET

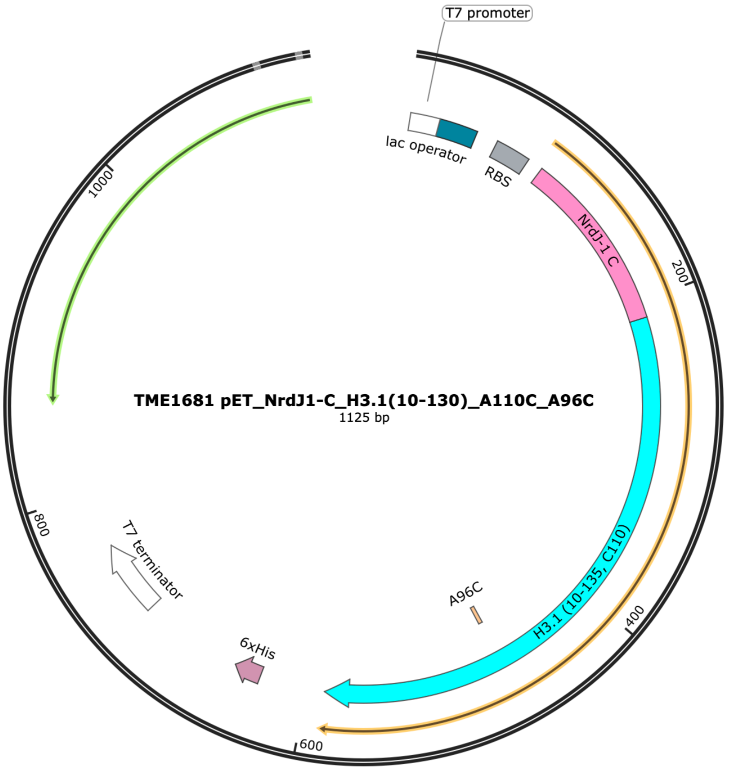


Expressed protein sequence (NrdJ-1^C-H3.1(10-130)):

MEAKTYIGKLKSRKIVSNEDTYDIQTSTHNFFANDILVHNSTGGKAPRKQLATKAAR
KSAPATGGVKKPHRYRPGTVALREIRRYQKSTELLIRKLPFQRLVREIAQDFKTDLRFQ
SSAVMALQEACEAYLVGLFEDTNLCAIHAKRVTIMPKDIQLARRIRGERA*

Flag-H3(1-28)-NrdJ-1$^{C}$-H3.1 in pCMV

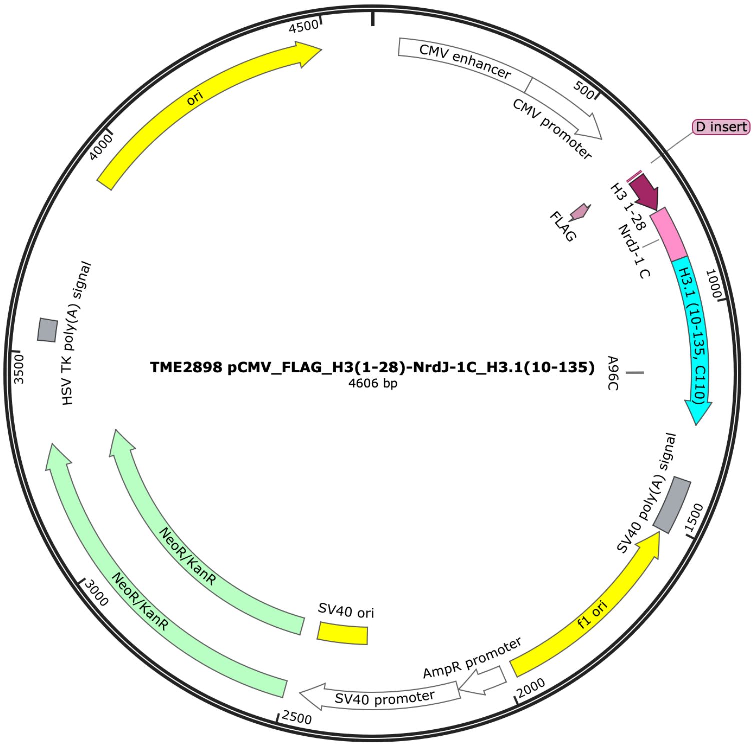


Expressed protein sequence (Flag-H3(1-28)-NrdJ-1$^{C}$-H3.1):

DYKDDDDKGARTKQTARKSTGGKAPRKQLATKAARKSMEAKTYIGKLKSRKIVSN
EDTYDIQTSTHNFFANDILVHNSTGGKAPRKQLATKAARKSAPATGGVKKPHRYRPG
TVALREIRRYQKSTELLIRKLPFQRLVREIAQDFKTDLRFQSSAVMALQEACEAYLVGL
FEDTNLCAIHAKRVTIMPKDIQLARRIRGERA*

<u>mCherry-LMNA-NrdJ-1$^{N}$ in pCMV</u>

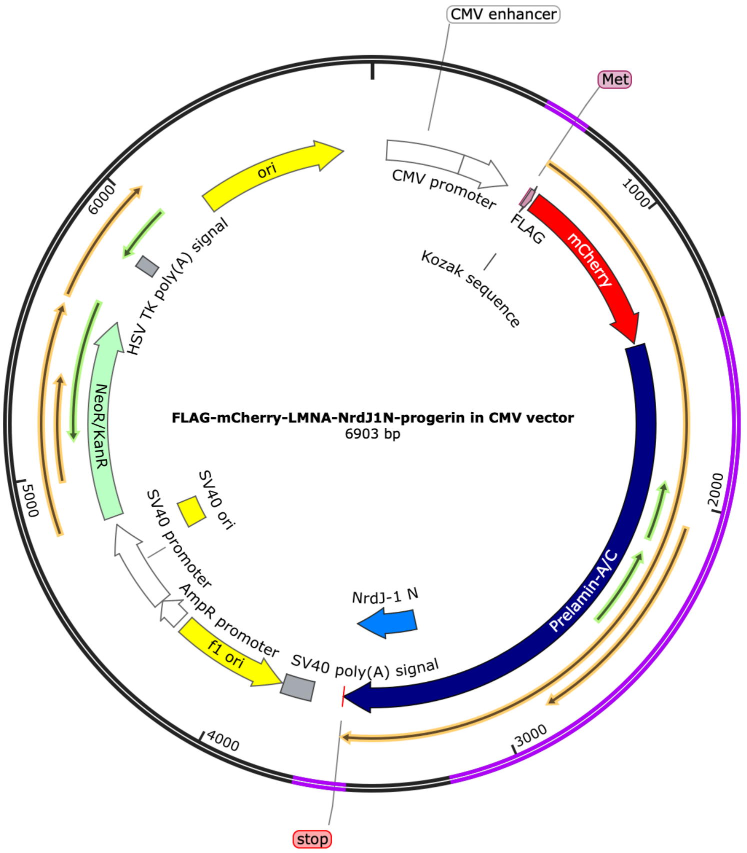


Expressed protein sequence (Flag-mCherry-progerin (NrdJ-1$^{N}$)):

MGDYKDDDDKGGSGMVSKGEEDNMAIIKEFMRFKVHMEGSVNGHEFEIEGEGEGR
PYEGTQTAKLKVTKGGPLPFAWDILSPQFMYGSKAYVKHPADIPDYLKLSFPEGFKW
ERVMNFEDGGVVTVTQDSSLQDGEFIYKVKLRGTNFPSDGPVMQKKTMGWEASSE
RMYPEDGALKGEIKQRLKLKDGGHYDAEVKTTYKAKKPVQLPGAYNVNIKLDITSH
NEDYTIVEQYERAEGRHSTGGMDELYKGGSGETPSQRRATRSGAQASSTPLSPTRITR
LQEKEDLQELNDRLAVYIDRVRSLETENAGLRLRITESEEVVSREVSGIKAAYEAELG
DARKTLDSVAKERARLQLELSKVREEFKELKARNTKKEGDLIAAQARLKDLEALLNS
KEAALSTALSEKRTLEGELHDLRGQVAKLEAALGEAKKQLQDEMLRRVDAENRLQT
MKEELDFQKNIYSEELRETKRRHETRLVEIDNGKQREFESRLADALQELRAQHEDQV
EQYKKELEKTYSAKLDNARQSAERNSNLVGAAHEELQQSRIRIDSLSAQLSQLQKQL
AAKEAKLRDLEDSLARERDTSRRLLAEKEREMAEMRARMQQQLDEYQELLDIKLAL
DMEIHAYRKLLEGEEERLRLSPSPTSQRSRGRASSHSSQTQGGGSVTKKRKLESTESR
SSFSQHARTSGRVAVEEVDEEGKFVRLRNKSNEDQSMGNWQIKRQNGDDPLLTYRFP
PKFTLKAGQVVTIWAAGAGATHSPPTDLVWKAQNTWGCGNSLRTALINSTGEEVAM
RKLVRSVTVVEDDEDEDGDDLLHHHHGSHCSSSGDPAEYNLRSRTVLCGTCGQPAD
KASASGCLVGSSEIITRNYGKTTIKEVVEIFDNDKNIQVLAFNTHTDNIEWAPIKAAQL
TRPNAELVELEIDTLHGVKTIRVTPDHPVYTKNRGYVRADELTDDDELVVAISGAQSP
QNCSIM*

Semisynthesis of H3 (1-9)-NrdJ-1$^{N}$ (C76A):

K(biotin)ARTKQ(ser)TARK(ac/me2/me3)CLVGSSEIITRNYGKTTIKEVVEIFDNDKNIQV
LAFNTHTDNIEWAPIKAAQLTRPNAELVELEIDTLHGVKTIRATPDHPVYTKNRGYVR
ADELTDDDELVVAI

(Underlined: modified or mutated residues during the studies)
